# The genetic origins of Saint Helena’s liberated Africans

**DOI:** 10.1101/787515

**Authors:** Marcela Sandoval-Velasco, Anuradha Jagadeesan, María C. Ávila-Arcos, Shyam Gopalakrishnan, Jazmín Ramos-Madrigal, J. Víctor Moreno-Mayar, Gabriel Renaud, Diana I. Cruz-Dávalos, Erna Johannesdóttir, Judy Watson, Kate Robson-Brown, Andrew Pearson, Agnar Helgason, M. Thomas P. Gilbert, Hannes Schroeder

**Affiliations:** Section for Evolutionary Genomics, The Globe Institute, Faculty of Health, University of Copenhagen, 1353 Copenhagen, Denmark; deCODE Genetics/Amgen, 101 Reykjavik, Iceland; Department of Anthropology, University of Iceland, 101 Reykjavik, Iceland; International Laboratory for Human Genome Research, National Autonomous University of Mexico, Juriquilla 76230, Santiago de Querétaro, Mexico; Lundbeck Centre for GeoGenetics, Department of Biology, University of Copenhagen, 1350 Copenhagen, Denmark; Department of Computational Biology, University of Lausanne, 1015 Lausanne, Switzerland; Swiss Institute of Bioinformatics, 1015 Lausanne, Switzerland; Department of Archaeology and Anthropology, University of Bristol, Bristol BS811, United Kingdom; Environmental Dimension Partnership, Atlantic Wharf, Cardiff CF10 4HF, UK; NTNU University Museum, Norwegian University of Science and Technology, 7491 Trondheim, Norway

## Abstract

Between the early 16^th^ and late 19^th^ centuries, an estimated 12 million Africans were transported to the Americas as part of the transatlantic slave trade. Following Britain’s abolition of slave trade in 1807, the Royal Navy patrolled the Atlantic and intercepted slave ships that continued to operate. During this period, the island of St Helena in the middle of the South Atlantic served as a depot for “liberated” Africans. Between 1840 and 1867, approximately 27,000 Africans were disembarked on the island. To investigate their origins, we generated genome-wide ancient DNA data for 20 individuals recovered from St Helena. The genetic data indicate that they came from West Central Africa, possibly the area of present-day Gabon and Angola. The data further suggest that they did not belong to a single population, confirming historical reports of cultural heterogeneity among the island’s African community. Our results shed new light on the origins of enslaved Africans during the final stages of the slave trade and illustrate how genetic data can be used to complement and validate existing historical sources.

## Introduction

The remote South Atlantic island of St Helena played an important role in the suppression of the transatlantic slave trade in the mid-19^th^ century. By that time the trade’s epicenter had shifted southwards from West Africa to West Central Africa (1). Strategically placed midway between the coasts of Angola and Brazil in the Atlantic Ocean, St Helena offered an ideal location for the Royal Naval vessels tasked with enforcing the Slave Trade Act. From 1840 onwards, the island was used as a depot for around 27,000 enslaved Africans who had been “recaptured” at sea (Fig.1, Table 1). It is estimated that around 8,000 of those people died on the island in the intervening years – most of whom were buried in two large graveyards in Rupert’s Valley (1) (Fig. S1).

**Figure 1.**
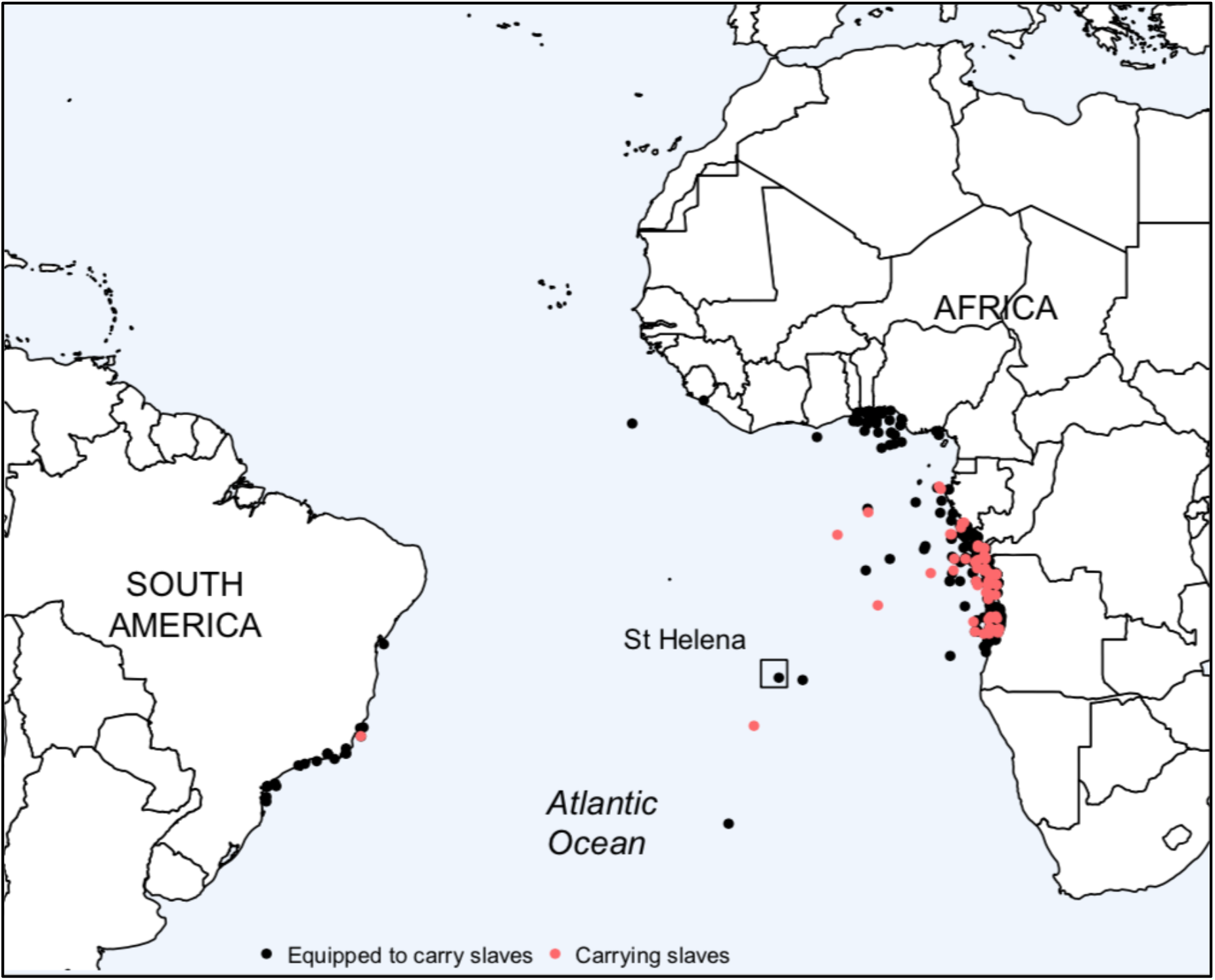
Location of ships captured by the Royal Navy. Locations of slave ships captured by the Royal British Navy between 1840 and 1867 and brought to St Helena according to ship logs and historical documents (1, 2).

**Table 1.** Number of prizes taken and Africans captured and disembarked on Saint Helena, 1840-1864 (1, 2).

| **Year of capture** | **Prizes taken** | **Slaves captured** | **Slaves liberated** |
| --- | --- | --- | --- |
| 1840-1844 | 33 | 6,552 | 5,852 |
| 1845-1849 | 25 | 9,313 | 8,333 |
| 1850-1854 | 11 | 3,953 | 3,350 |
| 1855-1859 | 5 | 1,970 | 1,694 |
| 1860-1864 | 13 | 5,206 | 4,988 |
| Total | 87 | 26,994 | 24,217 |

Archaeological excavations were conducted in Rupert’s Valley between 2007 and 2008 in connection with the construction of the island’s first airport, leading to the discovery of the skeletal remains of 325 individuals (2). The human remains and personal items recovered from some of the graves have yielded a wealth of information about the life and death of St Helena’s liberated African community. However, the graves provided few clues about their origins or cultural affiliations, beyond being African (1–3). Historical sources provide information about where the slave ships were captured (Fig. 1), but not about the origins of the “recaptured” slaves in Africa (1).

To investigate the origins of St Helena’s liberated Africans, we generated whole genome sequence (WGS) data (0.1-0.5× average coverage) from twenty individuals recovered during the archaeological excavations on the island and examined their genetic affinities to a set of 76 present-day African reference populations from 24 countries. Kinship analysis (4) revealed no close relationships between the sequenced individuals. However, chromosomal sex estimates (5) revealed many more male individuals (n=17) than females (n=3), in agreement with historical reports of a male bias in the transatlantic slave trade during its final phase (6).

## Results

Following initial screening of ancient DNA (aDNA) libraries from 63 individuals (Fig. S1 and Supplementary Table S1), we generated WGS data (0.1-0.5×) for a subset of twenty individuals (Table 2) using a whole genome capture method. Using this approach, we increased the human endogenous content of the aDNA libraries up to 10 times (Supplementary Table S2). All libraries exhibited features typical of authentic aDNA, including short average read lengths (<100 bp), characteristic fragmentation patterns, and an increased frequency of C-to-T substitutions at the 3′ and 5′ ends of reads (8) (Table 2, Supplementary Table S1 and S3). Contamination estimates were below 10% except for two samples: 254 and 351 (Table 2 and Supplementary Table S4). To explore the effect of contamination in those two samples, we reproduced a number of analyses using only damaged reads (7). As the results were qualitatively similar to those obtained when using the totality of reads, we used all mapped reads for the subsequent analysis from all the samples in the study.

**Table 2.** Sequencing statistics for the 20 individuals from St Helena.

| **Individual** | **Sex** | **DoC** | **%**<br>**C to T** | **mt**<br>**DoC** | **Contamination** | **mt hg** | **Ychr hg** |
| --- | --- | --- | --- | --- | --- | --- | --- |
| STH 213 | M | 0.5 | 4.8 | 168.1 | 0.6-1.3% | L1c3a | E1b1a1a1-M180 |
| STH 245 | M | 0.3 | 12.3 | 199.5 | 0.8-1.7% | L0a1b2a | E1b1a1a1-M180 |
| STH 248 | M | 0.3 | 8.6 | 179.1 | 1.5-2.5% | L0a1e | E1b1a1a1-M180 |
| STH 253 | M | 0.4 | 8.6 | 200.3 | 1.1-2.2% | L2a1f1 | E1b1a1a1-M180 |
| STH 254 | M | 0.1 | 5.7 | 28.9 | 35.6-52.8% | L3 | E1b1a1-M2 |
| STH 281 | M | 0.1 | 11.7 | 163.2 | 1.7- 3.6% | L3e1e | E1b1a1a1-M180 |
| STH 284 | F | 0.3 | 13.2 | 199.8 | 2.3-3.3% | L1b1a10b | - |
| STH 289 | M | 0.2 | 9.0 | 187.4 | 0.8-1.8% | L3e3b2 | E1b1a1a1-M180 |
| STH 344 | M | 0.2 | 11.1 | 187.4 | 0.1-1.2% | L3e1a3a | E1b1a1a1-M180 |
| STH 347 | F | 0.1 | 8.6 | 181.8 | 1.0-2.2% | L2a1f | - |
| STH 351 | M | 0.1 | 5.0 | 60.7 | 9.6-13.3% | L2b1a3 | E1b1a1a1-M180 |
| STH 358 | M | 0.2 | 15.7 | 198.0 | 1.3-2.6% | L3f1b1a | G2-L156 |
| STH 415 | M | 0.1 | 5.2 | 193.5 | 0.6-1.8% | L3d3a1 | E1b1a1a1-M180 |
| STH 436 | M | 0.1 | 8.1 | 189.5 | 0.5-2.0% | L3e2b1 | B2-M182 |
| STH 441 | M | 0.3 | 8.5 | 127.3 | 6.3-8.5% | L2a1f | E1b1a1a1-M180 |
| STH 460 | M | 0.2 | 13.9 | 199.6 | 0.1-1.3% | L3e1d1a | E1b1a1a1-M180 |
| STH 499 | M | 0.2 | 10.1 | 197.5 | 0.3-1.1% | L1b1a10 | E1b1a1a1-M180 |
| STH 514 | M | 0.2 | 12.3 | 197.0 | 0.5-1.4% | L2a1f1 | E1b1a1a1-M180 |
| STH 520 | M | 0.1 | 4.9 | 89.3 | 5.9-8.6% | L2b1a3 | E1b1a1a1-M180 |
| STH 524 | F | 0.3 | 11.9 | 65.6 | 6.7-10.4% | L2a1a3c | - |

Sex, genetic sex; DoC, average genomic depth of coverage; % C to T, observed frequency of C to T transitions at the first position of the 5' end of DNA fragments; mt DoC, mitochondrial DNA depth of coverage; Contamination, 95% CI for the mitochondrial DNA contamination estimates calculated using ContaMix (40); mt hg, mitochondrial DNA haplogroup, Ychr hg, Y-Chromosome haplogroup. For more information see Materials and Methods.

### Male bias in the transatlantic slave trade

Historical sources suggest that there was a male bias in the transatlantic slave trade, with two-thirds of enslaved individuals being male, and one-third female (6). Historical records also indicate that, as a result of various factors, these ratios varied widely over time and that during its last phase in the nineteenth century the trade was heavily biased towards adult males and children (6). To test if the genetic data fit with these observations, we determined the genetic sex of the individuals in our sample by calculating the proportion of reads mapping to the X and Y chromosomes, respectively (5). This revealed that 17 of the individuals were males and only three were females (Fig. S2 and Supplementary Table S5) – consistent with the strong bias reported in historical sources. For the 17 individuals deemed to be adults on the basis of skeletal morphology (Supplementary Table S6), genetic sex was in agreement with osteological inference. Three adolescents (12-18 years old) who could not be sexed based on skeletal traits, were confidently identified as males based on the genetic data. Among the 20 individuals under study, there were not any children. However, we note that out of the 325 individuals excavated, nearly half (177, 54.46%) were identified as subadults (younger than 18 years old) (2), also in agreement with an age bias reported in the historical records.

### Diverse origins in West Central Africa

To investigate the origins of the individuals from St Helena, we first determined their mitochondrial DNA (mtDNA) haplogroups and the Y-chromosome haplogroups for the males (7). While these markers only reflect a small portion of a person’s genetic ancestry, and are rarely able to provide definitive assignments of individual origins (9, 10), they can nonetheless be informative (11). We recovered complete mitochondrial genomes for all 20 individuals with an average depth of coverage between 30-200×, enabling confident assignment of mtDNA haplogroups (Table 2 and Supplementary Table S7). As expected, all individuals belonged to the L0-L3 clades of the African macrohaplogroup L. Based on the frequencies of subgroups from these clades across contemporary African populations (12, 13) we conclude that the individuals trace their matrilineal ancestry to West Central Africa (Fig. S4). In particular, the subgroups L2a1f, L2b, L1b1a, L1c3a, and L3f1b1a, found in 50% of the studied individuals, are commonly found in this region (13).

With regards to the Y chromosome, the vast majority (88%) of the 17 males under study belong to haplogroup E1b1a1a1-M180 (Fig. S5 and Supplementary Table S7). This haplogroup is found at high frequency in parts of West Central Africa today and has been associated with the Bantu expansion (14–17). Taken together, the mtDNA and Y chromosome evidence points at West Central Africa as the most likely place of origin of St Helena’s liberated Africans. However, due to the wide distribution of many of the mtDNA and Y chromosome lineages across the African continent as a result of the Bantu dispersals and other population movements (18, 19), it is difficult to identify more specific origins using uniparental markers alone.

To explore the origins of the individuals further, we performed a principal component analysis (PCA) using a reference panel consisting of 235,513 SNPs genotyped on 2,898 individuals from 76 African populations (Supplementary Table S8). The PCA was performed based on the overlapping SNPs by projecting the historical individuals onto the present-day variation (Supplementary Table S9). We find that the individuals from St Helena cluster with populations from West Central Africa, specifically with present-day populations from Cameroon, Gabon and Angola (Fig. 2A). These results are mirrored in the ADMIXTURE (20) analysis (K=7), which reveals that the African individuals have ancestry proportions that are most similar to those found in present-day Bantu-speaking populations from West Central Africa (Fig. 2B).

**Figure 2.**
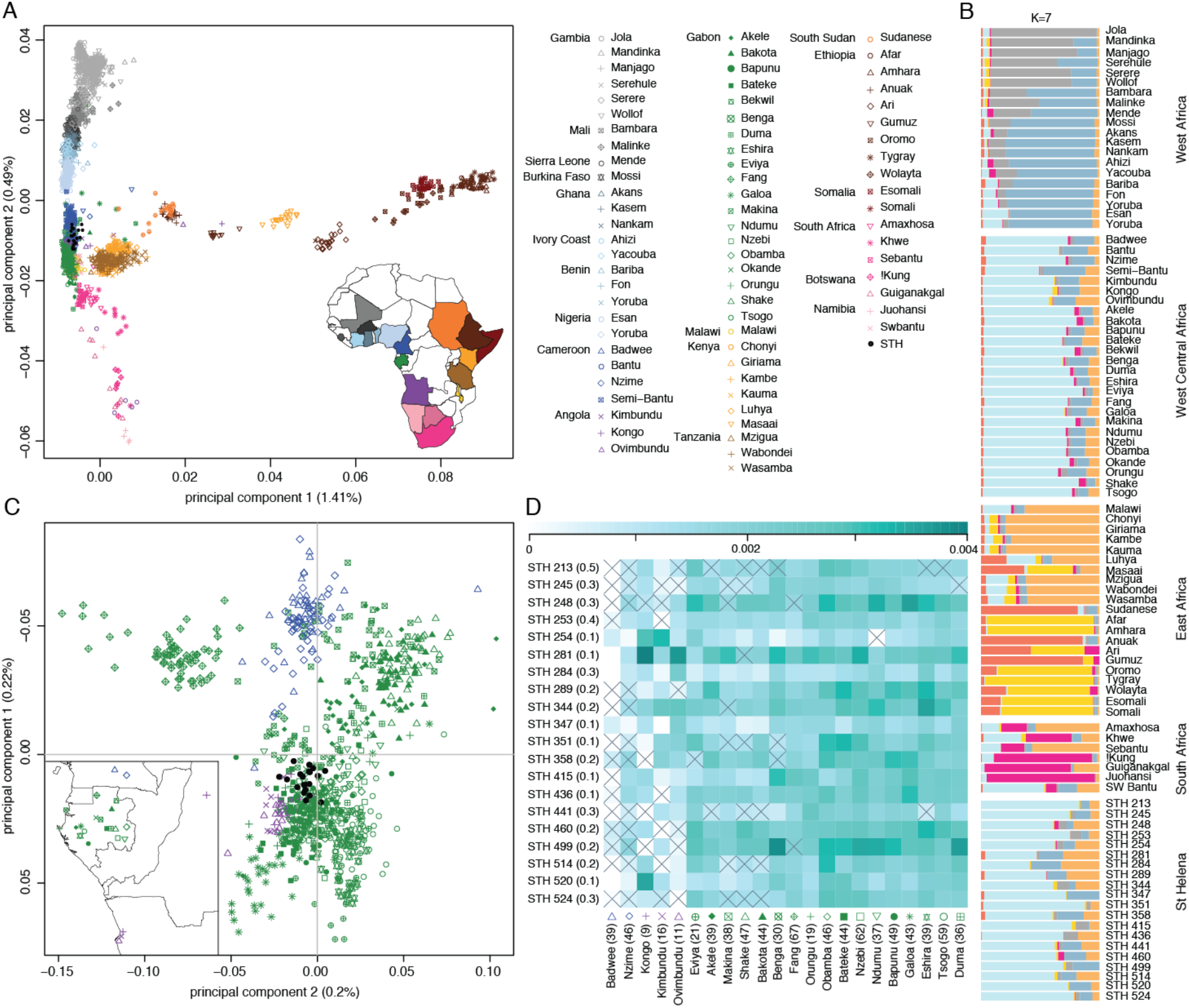
The genetic origins of Saint Helena’s liberated Africans. **(A)** principal components analysis with African reference populations from West, West Central, East and southern Africa; **(B)** ADMIXTURE analysis for K=7 showing population averages for reference populations; **(C)** principal components analysis showing only reference populations from West Central Africa; **(D)** heat map of mean pairwise identity-by-state scores weighted by allele frequency (wIBS). Populations that can be excluded as source populations based on a permutation test (7) are marked. The average depth of genomic coverage for each of the individuals in our sample and the number of individuals in the reference populations are given in parentheses.

Building on these findings, we then projected the individuals onto a PCA limited to 24 reference populations from Cameroon, Gabon and Angola and based on a total of 592,378 SNPs (Supplementary Table S10). In this analysis, the individuals from St Helena cluster more closely with present-day populations from Central Gabon and northern Angola than Cameroon (Fig. 2C). However, we note that while there clearly is genetic structure among these populations, we are unable to pinpoint specific source populations at this level of resolution. This problem is exacerbated by the fact that the actual source populations of the individuals under study might not be adequately represented in our refrence dataset.

To get a more direct handle on the genetic affinities of the African individuals and their origins, we computed a weighted measure of genomic identity by state (wIBS) (21) between each individual in our dataset and 841 individuals from 22 present-day reference populations from West Central Africa (7). Higher wIBS scores indicate stronger genetic affinity between the historical individual and any given reference population. The heatmap in Fig. 2D shows the mean wIBS scores for each of the individuals in our dataset (STH). To narrow down the number of potential source populations for each individual, we then performed a pairwise permutation test to compare the reference population that yielded the largest mean wIBS score with the mean wIBS scores obtained for the other 21 reference populations. This way, we were able to formally exclude up to 9 populations from the reference panel as possible sources for each of the individuals in our dataset (*P* <0.05/21, bonferroni corrected for 21 comparisons) (Fig. 2D and Fig. S10).

To further investigate and clarify these relationships, we computed a set of *D*-statistics of the form *D*(Outgroup, STH; refwIBS, test), where *refwIBS* is the population with the highest wIBS value for each individual from St Helena, and *test* represents all the other populations in the reference panel (7). These tests suggest that none of the tested populations are closer to the individuals in our dataset than the population with the maximum wIBS value, thus supporting the wIBS results (S12, Fig. S11 and Supplementary Table S11).

### Lack of close kinship ties supports diverse origins

In light of the shared mtDNA and Y chromosome haplogroups among some pairs of individuals, we explored whether any of them were close relatives. We used READ (4) that was developed specifically to infer kinship relationships in degraded samples, with as little as 0.1x shotgun coverage per genome for pairs of individuals. Exploring kinship relations among the individuals under study is important for two reasons. First, close kinship ties would suggest a common origin. Second, the presence of related individuals might shed new light on the dynamics of enslavement at the time. However, based on READ analysis we found all individuals to be unrelated. The lack of close kinship ties among the individuals in our sample supports the notion that the individuals had diverse origins.

## Discussion

Historical records offer important information about changes in the volume and routes of the transatlantic slave trade. In the case of St Helena, records from the island’s Vice Admiralty court indicate that the vast majority of slave ships brought to the island by the Royal Navy’s West Africa Squadron between 1840 and 1867 were captured south of the equator, with most captures clustered around the Central African coast (Fig. 1). However, while they give us a general indication where captives may have originated, these records reveal relatively little about their actual ethnic or geographic origins. Other historical records suggest that the captives arriving at St Helena were ethnically and linguistically diverse (1). Evidence from letters, diaries and official reports indicate in general terms that the Africans in St Helena’s depots were predominantly from Central and Southeast Africa, as this passage from a 19th century account of the island’s Liberated African Establishment illustrates: “With a few exceptions […] all the negroes who were brought to St. Helena came either from the countries bordering on the Congo River, from Angola, Benguela, or the Mozambique. The exceptions […] are natives of the interior of Africa” (22).

At the beginning of the 19th century there were three principal slaving regions in West Central Africa: northern Angola, Luanda and Benguela (23, 24). After 1807, international efforts to suppress the slave trade had a major impact on the operation of the trade in Luanda and Benguela, which had dominated the supply and shipment of slaves hitherto (24). This resulted in a shift and relocation of the trade to northern Angola, particularly the region between southern Gabon and Ambriz, passing by and making use of the Congo river to protect slavers from the British Navy and stretch the trade from the river banks all the way to the interior, expanding up to today’s Kinshasa (25). At this point, the slave trade from northern Angola was orders of magnitude bigger than in other regions, shipping approximately 340,000 captive Africans solely in the period between 1841-1867 (26, 27).

Our results are consistent with the historical evidence in that they suggest close affinities between the individuals recovered from Rupert’s Valley and present-day populations from West Central Africa. This is supported by the mtDNA evidence, as well as the Y-chromosome data and the genome-wide SNP-based analyses. While the mtDNA and Y-chromosome analyses lack in resolution, the genome-wide analyses suggest that the individuals had close affinities to present-day populations from Gabon and Angola. Based on the PCA, the ADMIXTURE analysis, as well as the patterns of wIBS, we further conclude that the individuals are likely to derive from several different populations and locations in West Central Africa, most probably within the boundaries of the northern Angola historical region, which at the time covered territories in today’s northern Angola, Congo, the DRC, and Gabon (25-27). In agreement with a multiple origin hypothesis, we find no genetic kinship relationships between the individuals under study.

Our ability to trace the genetic ancestry of enslaved individuals reliably is limited by the available reference panels, highlighting the need for more extensive sampling of human populations across the African continent, as currently there is relatively sparse information for most regions (28). In particular, the reference populations available from West Central Africa have a disproportionate number of individuals and populations from Gabon, but relatively few individuals from Angola and none from the Congo, the DRC, and other important places for the scope of this study. As larger reference samples from more populations become available, more accurate inferences regarding the origins of St Helena’s “liberated” Africans will be feasible. Furthermore, resolution is limited by the relatively small number of SNPs typically used in genotyping. Both of these problems are likely to be overcome in the near future with ever-increasing studies about African variation and the use of larger SNP panels and whole-genome sequencing (29).

In addition, the large-scale and widespread movements of people within the African continent itself during the period at which the slave trade was in operation, complicates the resolution of results. In this regard, it is important to note that the genetic and cultural landscape of sub-Saharan Africa has changed quite dramatically over the past 400 years (30). Therefore, we should be careful when applying present-day labels to past populations.

Our study highlights both the potential and challenges associated with the use of genetic data to shed light on the history of individuals in the transatlantic slave trade. Despite the limitations, we were able to confirm the region of origin for the individuals in our sample to West Central Africa and to narrow down the area between Gabon and Angola as the most likely geographical origin of the groups that these individuals belonged to. These findings are consistent with the limited historical records on St Helena that mention the Congo and Angola in this context. Moreover, they are consistent with historical research indicating a geographical shift during the latter stages of the transatlantic slave trade (6).

The individuals studied here represent only 20 among the more than 12 million enslaved Africans who were transported across the Atlantic between the early 16th and late 19th centuries. As there are many unanswered questions left, genomic approaches are proving to be powerful and accurate to validate and complement historical data, and to answer long-standing historical questions. In the future, the generation of more ancient genomic data, combined with adequate reference datasets and in-depth historical and ethnographic research, will contribute to our knowledge about the origins and movements of different peoples and groups in those historical times, offering the possibility of investigating personal and individual life histories (31). Despite not being representative of the entirety of people enslaved in transatlantic trade, the ability to tell the story of a few will help us to illustrate the condition of the many.

## Materials and Methods

### Samples and archaeological background

Premolar teeth were sampled from 63 individuals for ancient DNA (aDNA) analysis. Thirty-two of these were osteologically classified as males, 16 as female and 15 of unknown sex (2). Thirty individuals showed evidence of dental modification of various types, and all except 10 of them were classified as adults older than 18 years old (Supplementary Table S6).

To assess the level of aDNA preservation and the potential for genome-scale analysis we initially screened the samples using an Illumina-based shotgun sequencing approach. Most samples exhibited poor DNA preservation, with endogenous DNA contents ranging from 0.003% to 48% (mean 2.97%) (Supplementary Table S1). We therefore decided to apply an in-solution whole-genome DNA enrichment approach to a subset of 35 of the samples (Supplementary Table S2), which once sequenced yielded genome-wide data for 20 samples at a coverage ranging from 0.1× to 0.5×. We conducted a downsampling experiment to show that the data recovered was sufficient for population assignment against available reference data generated from present-day African populations (Fig. S6). For additional details see Supplementary Information.

### DNA extraction, NGS library preparation and sequencing

All experiments from sample processing, DNA extraction and library preparation (pre-library amplification) were conducted in dedicated aDNA clean lab facilities at the Centre for GeoGenetics, Natural History Museum of Denmark, University of Copenhagen, following established aDNA guidelines, including wearing coveralls, mask, hair and shoe covers, and double gloves; and cleaning working surfaces and all materials with bleach and 70% ethanol and subsequently irradiate them with ultraviolet light.

DNA was extracted from tooth roots using silica-based methods (32, 33). Recovered extract was then built into Illumina libraries using a blunt-end library preparation kit from NEB (E6070, New England Biolabs) and Illumina-specific adapters (34). Libraries were amplified and indexed, purified, and pooled for sequencing. Selected libraries were further enriched for the human fraction using the MYbaits Human Whole Genome Capture Kit (Arbor Biosciences). All sequencing was performed on an Illumina HiSeq 2500 platform run in 100bp single read chemistry mode.

### Data processing

After demultiplexing, adapter sequences were trimmed using AdapterRemoval 2.0 (35) with the default options in single-end mode. Adapter sequences, Ns, and low-quality tracks were trimmed from the reads and remaining reads shorter than 30 bp were removed. Reads were then mapped to the human reference genome (Hg19) using *bwa aln* algorithm (36) discarding duplicates and reads mapping to multiple regions of the genome. mapDamage 2.0 (37) was used to rescale the quality of bases that had a mismatch to the reference likely derived from damage. PMDtools (38) was used to filter BAM files and retain only reads showing signs of aDNA damage.

### Data authenticity

Authenticity of data was confirmed by: (a) sequencing of negative controls, (b) assessment of aDNA damage patterns and fragment length distribution, and (c) mtDNA contamination estimates. To examine damage parameters and validate sequencing data we used mapDamage 2.0 (37). We estimated contamination based on high coverage mtDNA data using two methods Schmutzi (39) and ContaMix (40).

### Sex determination

Genetic sex was determined by analyzing the ratio of X- to Y-chromosome reads following the study by Skoglund et al. (5) (Fig S2).

### Kinship analysis

To infer kinship relationships we used the software READ (4) and estimated relationships using a predefined panel of SNP sites consisting of 1,650,908 markers from the YRI individuals of the 1000 Genomes dataset.

### mtDNA analysis

Mitochondrial haplogroups were determined by retrieving the reads mapping to the rCRS (41) from the BAM files and creating a consensus sequence. Variants in the mtDNA were called using *SAMtools* and *bcftools* (42, 43), recorded in an hsd file and haplogroups were identified with the Haplogrep Software (44) using Phylotree build 17 as reference (45).

### Y chromosome DNA analysis

Inference of Y chromosome haplogroups for the male individuals was carried out using phylogenetically informative SNPs identified in studies of present-day Y chromosome diversity. We called Y-chromosome genotypes for each sample using the haploid genotype caller implemented in ANGSD (46), and performed a binary tree search to find the most derived SNP that determines the haplogroup of the individuals.

### Principal Component Analysis

PCA was performed using *smartpca* from the Eigensoft package (47) using the three different present-day datasets: (1) global, (2) Africa and (3) West Africa as reference panels. The ancient samples were projected onto the present-day populations using the option *lsqproject*. For the two samples with higher contamination estimates, in order to assess if contamination was affecting the overall results, we compared the “contaminated” samples with two samples with higher depth of coverage and lower levels of contamination by projecting onto the PCA using only reads with signs of aDNA damage (Fig. S7). We found similar results between samples, and decided to use all mapped reads from all the samples. However we caution that the results for the two contaminated samples (254 and 351) should be interpreted with care.

### ADMIXTURE Analysis

To investigate the genetic ancestry composition of the St Helena individuals we used the maximum-likelihood algorithm ADMIXTURE (20). We ran ADMIXTURE using different numbers of clusters (Ks). We first determined the values of K that produced the lowest cross-validation (CV) error values. For the whole Africa reference panel, the three lowest CV error values were obtained using K=6, K=7, and K=8. We ran 100 replicates of ADMIXTURE for each K and kept the replicate with the highest log-likelihood.

### Weighted IBS

To estimate the affinity of the studied individuals to possible African source populations, we performed an analysis based on a pairwise identity-by-state weighted by allele frequencies (wIBS) (21) in the West Central Africa panel where sample sizes were greater than nine. In order to test whether the reference population with the greatest mean wIBS for each of the individuals (*refwIBS*) was significantly greater than the value observed for other reference populations, we performed population pairwise permutation tests. In this test, where *REF* is another reference population, the wIBS values between the individuals in our sample and each member of reference population *refwIBS* and *REF* were shuffled 100,000 times to obtain the null distribution for the difference in mean wIBS values with each of the individuals in our sample (*refwIBS*-*REF*), under the assumption of no difference between them. For a single test, each of the individuals in our sample was deemed to be more closely related to population *refwIBS* than to *REF*, when 5% or fewer values from the null distribution were greater than the observed (non-permuted) difference between *refwIBS* and *REF*. These P values were further Bonferroni adjusted for the number of tests performed. Importantly, even if the true source population of an individual is not represented among the reference populations in our analysis, it is still possible to exclude reference populations with significantly lower wIBS values than *refwIBS*.

### D-statistics

*D*-statistics of the form *D*(Outgroup, STH; *refwIBS*, *REF*), where *refwIBS* is the population with the highest wIBS value for each individual from St Helena (STH) and *REF* represents any other given population in the reference panel, were computed using AdmixTools (48) and a set of 22 present-day reference populations from West Central Africa.

## Acknowledgements

We thank the community on St Helena, the St Helena government, and the St Helena National Trust for enabling us to carry out this work and to shed light on the history of the island’s “liberated” African community. We also thank the staff at the Danish National High-Throughput Sequencing Centre for technical assistance and N. Wales for experimental expertise and input on early stages of the manuscript, F. Vieira, J.A. Samaniego and H. Jónsson for assistance during data analysis, and I. Dull for thoughtful and valuable input into the manuscript. The research was supported by the EUROTAST project, a Marie Skłodowska-Curie Actions Initial Training Network funded by the European Union under grant agreement no. FP7/2007-2013/290344.

## Author contributions

M.S.-V., A.H., M.T.P.G., and H.S., designed research; M.S.-V., performed the experiments; M.S.-V., A.J., M.C.Á.-A., S.G., J.R.-M., J.V.M.-M., G.R., and D.I.C.-D., analyzed the data; E.J., J.W., K.R.-B., and A.P., provided samples and historical and archaeological background information; M.S.-V., A.H., M.T.P.G., H.S., A.J., M.C.Á.-A., S.G., J.R.-M., and J.V.M.-M., interpreted the results; M.S.-V. and H.S., wrote the manuscript with input from J.V.M.-M., A.H., M.T.P.G., S.G. and the remaining authors.

## References

1. Pearson AF (2016) Distant Freedom: St Helena and the Abolition of the Slave Trade, 1840-1872 (Liverpool University Press).

2. Pearson AF, Jeffs B, Witkin A, MacQuarrie H (2011) Infernal traffic: Excavation of a liberated African graveyard in Rupert’s Valley, St. Helena (Council for British Archeology).

3. MacQuarrie H, Pearson A (2016) Prize Possessions: Transported Material Culture of the Post-Abolition Enslaved-New Evidence from St Helena. Slavery and Abolition 37(1):45–72.

4. Monroy Kuhn JM, Jakobsson M, Günther T (2018) Estimating genetic kin relationships in prehistoric populations. PLoS One 13(4):e0195491.

5. Skoglund P, Storå J, Götherström A (2013) Accurate sex identification of ancient human remains using DNA shotgun sequencing. Journal of Archaeological Science 40:4477–4482.

6. Eltis D, Engerman SL (1993) Fluctuations in sex and age ratios in the transatlantic slave trade, 1663-1864 1. Econ Hist Rev 46(2):308–323.

7. See Materials and Methods

8. Briggs AW, et al. (2007) Patterns of damage in genomic DNA sequences from a Neandertal. Proc Natl Acad Sci U S A 104(37):14616–14621.

9. Salas A, Carracedo Á, Richards M, Macaulay V (2005) Charting the ancestry of African Americans. Am J Hum Genet 77(4):676–680.

10. Ely B, Wilson JL, Jackson F, Jackson BA (2006) African-American mitochondrial DNAs often match mtDNAs found in multiple African ethnic groups. BMC Biol 4:34.

11. Schroeder H, et al. (2015) Genome-wide ancestry of 17th-century enslaved Africans from the Caribbean. Proc Natl Acad Sci U S A 112(12):3669–3673.

12. Salas A, et al. (2002) The making of the African mtDNA landscape. Am J Hum Genet 71(5):1082–1111.

13. Cerezo M, et al. (2016) Comprehensive Analysis of Pan-African Mitochondrial DNA Variation Provides New Insights into Continental Variation and Demography. J Genet Genomics 43(3):133–143.

14. Wood ET, et al. (2005) Contrasting patterns of Y chromosome and mtDNA variation in Africa: evidence for sex-biased demographic processes. Eur J Hum Genet 13(7):867–876.

15. Montano V, et al. (2011) The Bantu expansion revisited: a new analysis of Y chromosome variation in Central Western Africa. Mol Ecol 20(13):2693–2708.

16. Poznik GD, et al. (2016) Punctuated bursts in human male demography inferred from 1,244 worldwide Y-chromosome sequences. Nat Genet 48(6):593–599.

17. Cruciani F, et al. (2011) REPOR TA Revised Root for the Human Y Chromosomal Phylogenetic Tree: The Origin of Patrilineal Diversity in Africa. Am J Hum Genet 88(6):814–818.

18. Salas A, et al. (2002) The making of the African mtDNA landscape. Am J Hum Genet 71(5):1082–1111.

19. Beleza S, Gusmão L, Amorim A, Carracedo A, Salas A (2005) The genetic legacy of western Bantu migrations. Human Genetics 117(4):366–375.

20. Alexander DH, Novembre J, Lange K (2009) Fast model-based estimation of ancestry in unrelated individuals. Genome Res 19(9):1655–1664.

21. Jagadeesan A, et al. (2018) Reconstructing an African haploid genome from the 18th century. Nat Genet 50(2):199–205.

22. McHenry G (1845) An Account of the Liberated African Establishment on St. Helena. Simmond’s Colonial Magazine and Foreign Miscellany 5:172–183.

23. Miller J (1975) Legal Portuguese slaving from Angola. Some preliminary indications of volume and direction. Outre-Mers Revue d’histoire 62(226):135–176.

24. Ferreira R (2011) The suppression of the slave trade and slave departures from Angola, 1830s-1860s. História Unisinos 15(1):3.

25. Wissenbach MCC (2015) Dinâmicas históricas de um porto centro-africano: Ambriz e o Baixo Congo nos finais do tráfico atlântico de escravos (1840 a 1870). Revista de História (172):163–195.

26. da Silva DBD (2013) The Atlantic Slave Trade from Angola: A Port-by-Port Estimate of Slaves Embarked, 1701–1867. Int J Afr Hist Stud 46(1):105–122.

27. da Silva DBD (2017) The Atlantic Slave Trade from West Central Africa, 1780–1867 (Cambridge University Press).

28. Popejoy AB, Fullerton SM (2016) Genomics is failing on diversity. Nature 538(7624):161–164.

29. Gurdasani D, et al. (2015) The African Genome Variation Project shapes medical genetics in Africa. Nature 517(7534):327–332.

30. Lovejoy PE (1989) The Impact of the Atlantic Slave Trade on Africa: A Review of the Literature. The Journal of African History 30(3):365–394.

31. Schablitsky JM, et al. (2019) Ancient DNA analysis of a nineteenth century tobacco pipe from a Maryland slave quarter. Journal of Archaeological Science 105:11–18.

32. Dabney J, et al. (2013) Complete mitochondrial genome sequence of a Middle Pleistocene cave bear reconstructed from ultrashort DNA fragments. Proc Natl Acad Sci U S A 110(39):15758–15763.

33. Rohland N, Hofreiter M (2007) Ancient DNA extraction from bones and teeth. Nat Protoc 2(7):1756–1762.

34. Meyer M, Kircher M (2010) Illumina Sequencing Library Preparation for Highly Multiplexed Target Capture and Sequencing. Cold Spring Harbor Protocols 2010(6):db.prot5448–pdb.prot5448.

35. Schubert M, Lindgreen S, Orlando L (2016) AdapterRemoval v2: rapid adapter trimming, identification, and read merging. BMC Res Notes:1–7.

36. Li H, Durbin R (2009) Fast and accurate short read alignment with Burrows–Wheeler transform. Bioinformatics 25(14):1754–1760.

37. Jonsson H, Ginolhac A, Schubert M, Johnson PLF, Orlando L (2013) mapDamage2.0: fast approximate Bayesian estimates of ancient DNA damage parameters. Bioinformatics 29(13):1682–1684.

38. Skoglund P, et al. (2014) Separating endogenous ancient DNA from modern day contamination in a Siberian Neandertal. Proc Natl Acad Sci U S A 111(6):2229–2234.

39. Renaud G, Slon V, Duggan AT, Kelso J (2015) Schmutzi: estimation of contamination and endogenous mitochondrial consensus calling for ancient DNA. Genome Biol 16:224.

40. Fu Q, et al. (2013) A revised timescale for human evolution based on ancient mitochondrial genomes. Curr Biol 23(7):553–559.

41. Andrews RM, et al. (1999) Reanalysis and revision of the Cambridge reference sequence for human mitochondrial DNA. Nat Genet 23(2):147.

42. Li H, et al. (2009) The Sequence Alignment/Map format and SAMtools. Bioinformatics 25(16):2078–2079.

43. Li H (2011) A statistical framework for SNP calling, mutation discovery, association mapping and population genetical parameter estimation from sequencing data. Bioinformatics 27(21):2987–2993.

44. Kloss-Brandstätter A, et al. (2011) HaploGrep: a fast and reliable algorithm for automatic classification of mitochondrial DNA haplogroups. Hum Mutat 32(1):25–32.

45. van Oven M, Kayser M (2009) Updated comprehensive phylogenetic tree of global human mitochondrial DNA variation. Hum Mutat 30(2):E386–94.

46. Cruz-Dávalos DI, et al. (2018) In-solution Y-chromosome capture-enrichment on ancient DNA libraries. BMC Genomics 19(1). doi:10.1186/s12864-018-4945-x.

47. Patterson N, Price AL, Reich D (2006) Population Structure and Eigenanalysis. PLoS Genetics 2(12):e190.

48. Patterson N, et al. (2012) Ancient admixture in human history. Genetics 192(3):1065–1093.

