## Supplementary Information and Figures for "The genetic origins of Saint Helena’s liberated Africans"

**This PDF includes:**

Supplementary Information
Supplementary Figures S1 to 11
References

**Other supplementary materials for this manuscript include:**

Datasets S1 to S11

### Supplementary Information

#### Historical and archaeological background

Following its discovery in 1502 by the Portuguese, St Helena became a maritime staging post, benefitting from its strategic location in the middle of the South Atlantic Ocean between the coast of Angola and Brazil. In 1659 the island became a British colony, and apart from the brief occupation by the Dutch in 1673, it has remained a British territory to the present day.

This remote island played only an indirect role in the suppression of the slave trade during the initial decades after the passing of Britain's abolition act in 1807, during which time Sierra Leone was the most significant naval outpost for the British anti-slavery campaign on the African coast and in the Atlantic. It was only after 1839 that the role of St Helena in the slave trade changed dramatically as a consequence of the Slave Trade (Portugal) Act, which authorized British warships to search and detain Portuguese vessels equipped for the slave trade, and for vice-admiralty courts to judge and condemn them. Critically too, the Act permitted warships to patrol south of the equator, and thus to police the previously immune slaving zones on the West-Central African coast.

In 1839 a vice-admiralty court was constituted in St Helena and a camp (or 'depot') for liberated Africans was established. The first cohorts of liberated Africans were accommodated in Lemon Valley (located 3km to the south-west of Jamestown, the island's capital). However, this site presented numerous difficulties, and consequently the depot was relocated to Rupert's Valley, which offered better conditions for accommodating the camp and could host a larger number of people. The flux of African "recaptives" into St Helena varied enormously. At times the depot held over a thousand people, while at other periods it would be almost empty. The apogee of St Helena's role in the anti-slavery movement started in 1845 when the Brazilian slave trade came under increasing military and diplomatic pressure; this included the passage of a new parliamentary act which authorized British cruisers to capture Brazilian slavers both north and south of the equator and to bring them for adjudication. St Helena was ideally placed to receive captured slave ships and their human cargo, and in total the island would receive around 27,000 captives between the opening of its depots in 1840 and the collapse of the Atlantic Slave Trade in the mid-1860s.

Historical records report that the captives arriving at St Helena were ethnically and linguistically diverse. Evidence from letters, diaries and official reports indicate in general terms that the Africans in St Helena's depots were predominantly from Central and Southeast Africa, sometimes with the perception of an interior rather than a coastal group of people. Furthermore, data from the vice-admiralty court shows that all slave-laden prizes directed to St Helena came from south of the equator, with most captures clustered around the Central African coast (1).

Between 2007 and 2008, archaeological excavations took place in St Helena as part of a mitigation programme for the construction of the first airport on the island. Human remains of 325 individuals were recovered from a total of 178 graves (Fig. S1), excavated from one of two large unmarked graveyards in Rupert's Valley (2). The samples in this study were collected in 2012 from the 'Liberated African Burial' skeletal collection located in the Museum of Saint Helena. We received premolar teeth samples from 63 individuals with various degrees of preservation and skeletal completeness, of which 32 were osteologically classified as males, 16 as female and 15 of unknown sex (2). Thirty individuals showed evidence of dental modification of various types, and all except 10 of them were classified as adults older than 18 years old (Table S6).

### Sample processing, DNA extraction and library preparation

Sixty-three premolar teeth samples were initially processed and screened for ancient DNA (aDNA) content and preservation. The surface of the samples was first wiped with a tissue dipped in 10% bleach solution. Subsequently, to remove any adhering sediment and to minimize potential contamination from modern DNA due to previous handling, the surface layer of the tooth was removed using the edge of a cutting disk fitted in a mechanical drill. Samples were then UV-irradiated for 2 min on each side to further reduce the amount of surface contaminants. Afterwards, the crown of the tooth was cut off from the root using again a cutting disk, and the tooth root was ground to powder using a ball mill and the Micro-Dismembrator S (Braun Biotech, Allentown, PA). Between 100-300 mg of root powder was then digested overnight at 45ºC in 5 ml digestion buffer containing 4.7 ml 0.5 M EDTA, 50 µL Proteinase K (0.14-0.22 mg/ml, Roche, Basel, CH) and 250 µL 10% N-Laurylsarcosyl. Following incubation, two silica-based purification methods (3, 4) were used to isolate the DNA from the supernatant. Extractions were carried out in batches of 8 samples each with extraction blanks in order to monitor cross-contamination and contamination from reagents. A total of 124 DNA extracts were recovered by elution of the DNA in 60µL of EB buffer (Qiagen, Valencia, CA).

Double stranded libraries were prepared using a blunt-end library preparation kit from NEB (E6070) (New England Biolabs, Ipswich, MA) and Illumina specific adapters (5) according to the manufacturer's instructions, with the following minor modifications. We skipped the initial nebulization step because of the fragmented nature of aDNA. End-repair was performed in 50 µl reactions and for each of the samples 42.5 µL of the aDNA extract was used as starting template. For the end-repair reaction, samples were incubated in a thermocycler for 20 minutes at 12°C followed by 15 minutes at 37°C. During the adaptor ligation step samples were incubated in a thermocycler for 20 minutes at 20°C. No size selection with magnetic beads was performed, and the fill-in reaction was incubated for 20 mins at 65°C followed by 20 mins at 80°C to inactivate the Bst enzyme.

Samples were indexed and amplified in 100 µl PCR reactions, containing 20 µl of aDNA library template, 10 µl 10X PCR buffer, 10 µl MgCl2 (25 mM), 0.8 µl BSA (20 mg/ml), 0.8 µl dNTPs (25 mM), 2 µl of each primer (10 µM, inPE forward primer and indexed reverse primer), and 0.8 µl AmpliTaq Gold DNA Polymerase (Applied Biosystems, Foster City, CA). Thermocycling conditions were 5 min at 95°C, followed by 10-14 cycles of 30s at 95°C, 30s at 60°C and 40s at 72°C, and a final 7 min elongation step at 72°C. The number of cycles was estimated for each sample using qPCR. Following amplification, samples were purified using Qiaquick columns (Qiagen) according to manufacturers' instructions and quantified either using a 2200 TapeStation Instrument or an Agilent 2100 BioAnalyzer (Agilent Technologies, Palo Alto, CA, USA). Samples were then pooled in equimolar amounts and screened via shotgun sequencing across 3 lanes of an Illumina HiSeq 2500 platform run in 100bp single read chemistry mode.

### Whole genome enrichment

Based on shotgun sequencing screening results, which showed varying amounts of endogenous DNA between the samples (0.003 – 48%) (Table S1), thirty-five samples with a wide range of human DNA content (0.1%-21.6%) were selected for whole-genome capture (WGC) experiments and further sequencing in order to enrich for the human portion in the samples and increase informative data output (Table S2). We excluded the sample with 48% endogenous content and sequence that one deeper without further processing. For enrichment, we used the Mybaits Human Whole Genome Capture Kit (Arbor Biosciences, Ann Arbor, MI) that uses biotinylated RNA probes transcribed from genomic DNA libraries to capture the human DNA in the aDNA libraries. We followed manufacturer's instructions as in the Version 1.3.8 of the Mybaits user manual but we modified it slightly, following instructions taken from Version 2.3.1 (https://arborbiosci.com/mybaits-manual/), particularly instead of releasing the captured DNA target molecules from the RNA baits using a NaOH treatment (as suggested in version 1.3.8) we resuspended the beads with 30 µl Molecular Biology Grade Water, and followed with PCR directly after finishing the capture.

After capture, all samples were amplified for 14 cycles in 50 µl PCR reactions, containing 15 µl of aDNA library as template, 25 µl 2X KAPA HotStart ReadyMix, 1 µl µl BSA (20 mg/ml), 1.5 µl of 10 µM agnostic primers IS5 and IS6 (Meyer and Kircher, 2010), and 7 µl of Molecular Biology grade water Thermocycling conditions were 3 min at 98°C, followed by 14 cycles of 20 s at 98°C, 30 s at 60°C and 30 s at 72°C, and a final 5 min elongation step at 72°C.

Following amplification, the captured libraries were purified using Qiaquick columns (Qiagen) according to manufacturer's instructions and quantified using a 2200 TapeStation Instrument (Agilent Technologies), pooled in equimolar amounts, and sequenced on 3 lanes of an Illumina HiSeq 2500 run in 100 SR mode.

### Sequencing and data processing

Based on capture efficiency results, we decided to deep sequence samples fulfilling the following requirements: 1) samples should have more than 5% endogenous content after WGC, 2) samples should have at least 0.05 depth of coverage after WGC, and 3) samples should present an average of at least 10% of damage nucleotides along the reads. Following these cut-off, we ended up choosing 20 samples (Table 2) that we sequenced deeper on 3 lanes of an Illumina HiSeq 2500 under the 100bp single read mode.

Raw sequencing data was basecalled using Illumina software CASAVA 1.8.2 and sequences were demultiplexed according to the 6 nucleotide index that was used for library preparation for each sample. Using *AdapterRemoval v2* (6) with the default options in single-end mode, adapter sequences, Ns, and low-quality tracks were trimmed from the reads and remaining reads shorter than 30 bp were removed. Using *bwa aln* algorithm (7), trimmed reads were individually aligned and mapped to the human reference genome build 37 (hg19), including the revised Cambridge Reference Sequence (rCRS) for the mitochondrial genome (8). The resulting alignments were filtered for reads with a mapping quality below 30, and duplicate reads using *SAMtools* rmdup function (9) and *AWK* commands were used to remove reads having alternative mapping coordinates by controlling for XA, XT and X0 tags. BAM files for every sample sequenced more than once were merged with *samtools merge* and duplicates were removed again after the merging step with *SAMtools rmdup*. The quality of the aligned bases was rescaled with mapDamage2 (10).

Once duplicates and ambiguously mapped reads were removed, quality and features of the libraries were estimated. Unique endogenous fractions were calculated by dividing the number of uniquely mapped reads (reads remaining after removing duplicates) by the number of reads after adapter removal and QC (raw reads). Average depth of coverage was calculated by dividing the number of uniquely mapped reads times the average read length by the size of the human reference genome used (hg19), that excluding Ns, contains 2.86 Gb.

### MapDamage analysis

Ancient DNA is characterized by patterns of molecular damage caused by hydrolytic and oxidative processes. These chemical reactions severely degrade the DNA backbone, making it prone to fragmentation, and introduce a series of chemical modifications to nucleotide bases. The most prominent consists on the loss of a methyl group in the cytosines, called deamination. During PCR, deamination can result in a nucleotide misincorporation in which cytosine is misidentified by the DNA polymerase as a thymine, and copying of the damaged molecule also leads to replacement of guanine by adenine on the complementary strand, giving a characteristic C-to-T and G-to-A transitions at the end of the 5' and 3' fragments respectively. Furthermore, ancient DNA is expected to display long single stranded overhangs (11).

To validate sequencing data based on these aDNA characteristics we used mapDamage 2.0 (10) and examined three damage parameters for each sample: 1) the frequency of C-to-T transitions in the first position at the 5' end of reads, 2) λ, the fraction of bases positioned in single-stranded overhangs, and 3) δs, the estimated C-to-T transition rate in the single-stranded overhangs (Table S3). As expected, sequencing data presented short average read lengths ranging from 43 to 86 bp. Furthermore, PMDtools (12) was used to filter bam files and retain only reads showing signs of aDNA damage.

**Contamination estimates**

We estimated contamination based on high coverage mtDNA data using two methods Schmutzi (13) and ContaMix (14) (Table S4). Schmutzi was used to jointly infer the endogenous consensus and to estimate present-day human contamination levels. The program was run twice, once with the "–notusepredC" option to use potential contaminant mitogenomes in the database provided with the software and once without which allows the predicted contaminant mitogenome to be included in this database. In both instances, the "–uselength" option was used to utilize the length of the molecules in addition to misincorporation patterns due to deamination to distinguish the endogenous base from the contaminant one. The predicted endogenous bases and insertions/deletions are produced with an error probability. The endogenous consensus sequences were filtered using a quality filter of 50 on a PHRED scale which represent an error probability of 1 in 100,000.

ContaMix estimates the proportion of contamination in sequences aligning to the human mtDNA with a likelihood-based method using a Markov chain Monte Carlo probabilistic model to estimate the contamination rate. The method generates a moment-based estimate of the error rate and a Bayesian-based estimate of the posterior probability of the contamination fraction. After mapping the reads for each sample to the nuclear genome and the rCRS, we extracted those mapping only to the rCRS and determined a mtDNA consensus sequence using a simple majority rule. We then re-mapped the extracted reads, but now to the consensus sequence that was created, and aligned the consensus to a set of 311 mtDNA sequences from around the world (15) as done in Fu et al. (14). We used the 311 individuals alignment and the re-mapped reads as input for the contamination estimate algorithm ContaMix. We ran three chains of 50,000 iterations for the Monte Carlo Markov Chain and discarded the first 10,000.

**Genetic sex determination and kinship analysis**

The ratio of reads mapping to Y and the X chromosome was used to determine the sex of each sample as described in Skoglund et al. (16). Discarding alignments with mapping quality lower than 30, we calculated the fraction of reads mapping to the Y chromosome out of the total of reads mapping to both the Y and the X chromosome which in turn was used to assign the sample to either XX or XY (Table S5).

To investigate the kinship relationships between the STH individuals we used the software READ (17) which was developed specifically for whole genome shotgun sequencing of ancient samples. READ works by partitioning the genome in non-overlapping windows of 1 Mbps and calculates the proportions of haploid mismatches and matches, P0 and P1, for each window. Since P0+P1 = 1, READ uses P0 as a single test statistic. READ defines three thresholds to identify pairwise relatedness as unrelated, second-degree (i.e. nephew/niece-uncle/aunt, grandparent- grandchild or half-siblings), first-degree (parent-offspring or siblings) and identical individuals/identical twins. There are four possible outcomes when running READ: unrelated (normalized P0≥0.9), second degree (0.9≥normalized P0≥0.8), first degree (0.8≥normalized P0≥0.65) and identical twins/identical individuals (normalized P0<0.65). Kinship values were estimated using a predefined panel of SNP sites consisting of 1,650,908 markers taken from the YRI individuals of the 1000 Genomes dataset.

**Mitochondrial DNA and Y-chromosome analysis**

To call Y-chromosome genotypes for each sample, we used the haploid genotype caller implemented in ANGSD, by sampling a random base at each position of the Y-chromosome and retaining only bases with quality scores of at least 13 (18). Next, we performed a binary tree search to find the most derived SNP that determines the haplogroup of the individuals as done in Cruz-Dávalos et al. (19). We used as input the phylogenetic tree constructed from the Y-SNPs reported in Phase 3 of the 1000 Human Genomes Project (20).

**Reference datasets**

As a global reference dataset we used the public reference panel HGDP (21) as distributed in Malaspinas et al. (22). The selected panel includes 644,117 SNPs, genotyped on 938 individuals from 53 populations spanning 6 geographic regions around the world (Africa, East Asia, West-Central-South Asia, Europe, Oceania and America) (Fig S3).

For the Africa reference panel, we merged publicly available genotype data from several studies and put together a dataset consisting of 235,513 SNPs from 2898 individuals from 76 different African populations spanning West West Central, South and East African regions (Table S8).

For the zoom-in West Central African (WCA) dataset, we merged two recently generated datasets. For the first dataset, we extracted 73 individuals from the two non-hunter-gatherer populations (Nzime from Cameroon and Nzebi from Gabon) present in the publicly available dataset published by Patin et al. (23) consisting of 930,134 markers genotyped on the Illumina Omni1 chip. We merged them with a second dataset containing the non-hunter-gatherer populations from the study published by Patin et al. (24) consisting of 957 individuals and 690,739 autosomal markers genotyped on the Illumina 1M chip. Before merging we filtered out monomorphic sites and kept only biallelic SNPs in each of the datasets. After excluding also multiallelic markers in both datasets, the biallelic datasets were merged resulting in an overlapping marker set of 658,726 SNPs. A total of 53 individuals were used from dataset one (23) and 920 individuals were used from dataset two (24). After merging them, we removed markers where more than than 10% of individuals of a population are missing genotypes for a given marker, with which we removed 30,192 markers. We then applied *smartpca* (Eigensoft v6.1.3) (25) on the panel using LD regression and allowed smartpca to remove outliers. This generated the final dataset consisting of 592,378 SNPs and 851 individuals from 24 populations listed on Table S10.

**Principal component analysis**

In order to merge the reference panel with the ancient individuals' genome-wide data and perform Principal Component Analysis (PCA), we randomly sampled one allele at each site and for each individual in the reference panel and made such site homozygous for the drawn allele, as described in Skoglund et al. (26).

Using *SAMtools (9) mpileup*, we retrieved the alignment information from the bam files of the ancient samples only at the sites present in the genotype reference file. If covered, most sites had a single read, so if the queried base had a quality above 20, then such base was considered and reported as homozygous at that site. For the few cases where more than one read covered the queried site, a random allele was selected from those with base quality above 20, and again it was reported as homozygous at that site. The pseudo-haploid genotypes were then merged with the present-day reference panels using PLINK (27). This resulted in merged datasets with varying number of overlapping SNPs between the reference and the ancient sample, as shown in Table S9 for the different reference datasets. To perform PCA, we ran *smartpca* (Eigensoft v6.1.3) (25) using each merged ancient-reference panel genotype file prepared as described above. We also ran *smartpca* only on the diploid reference panel to calculate the reference-only eigenvectors. Eigenvectors were plotted independently for each dataset using R (http://www.r-project.org/). In order to visualize the 20 ancient samples in a single PCA scatter plot we used two projection approaches, the *lsqproject* option of *smartpca* and a Procrustes transformation using the R VEGAN package (http://vegan.r-forge.r-project.org) as done in Skoglund et al. (26).

To investigate if the low coverage data recovered was sufficient for population assignment against available reference panels, we conducted a downsampling experiment of overlapping sites in which we took the sample with the highest number of overlapping sites (STH 213 with 72,587 sites) and randomly downsample the number of sites to 60k, 30k and 15k with 10 iterations each time. We then performed Principal Component Analysis to show that the decreasing number of overlapping sites didn't really affect the positioning of the sample in the PCA space in relation to the reference panel, and hence the population assignment (Fig. S6).

### ADMIXTURE analysis

We ran unsupervised ADMIXTURE (28) on the whole Africa dataset using default parameters for K = 2 to K = 10 (Fig. S8 and S9). We ran 10 replicates for each K and based on CV error values we picked the Ks with the lowest values. The lowest CV error values were obtained using K=6, K=7, and K=8. We ran a total of 100 replicates for these Ks, selecting K=7 and choosing the replicate with the highest log-likelihood as result for that K.

### Weighted IBS (wIBS) analysis

In the weighted IBS (wIBS) method (29), alleles are weighted according to their frequencies. Diploid genotypes from reference African individuals are used when calculating wIBS, and missing genotype loci are ignored. For example if *a0* and *a1* are two alleles at a locus, weight for *a0* can be defined as:

$$Wt_{a0} = \frac{1 - Frq_{a0^2}}{Frq_{a0} \times Frq_{a1}}$$

A weight for *a1* can also be similarly defined. The weighted IBS score between two individuals *p* and *q,* across *N* overlapping SNPs, ignoring loci where either p or q have a missing genotype, is calculated as:

$$wIBS(p,q) = \sum_{i=0}^{N} \frac{No.of\ Shared\ Copy\ (a0) \times No.of\ Shared\ Copy\ (a1) \times Wt_{a1}}{2 \times Max\ (Wt_{a0}\ , Wt_{a1})}$$

This has an effect of up-weighting the IBS scores when rare alleles are shared.

We calculated wIBS scores for all ancient individuals against the West Central Africa dataset, excluding the individuals from populations of Bekwil and Okande from Gabon as sample sizes were smaller than 9 individuals.

In order to test whether the population with the greatest mean wIBS for each of the individuals (A) was significantly greater than the value observed for other reference populations, we performed population pairwise permutation tests. In this test, where B is another reference population, the wIBS values between the individuals in our sample and labels of the members of reference population A and B were shuffled 100,000 times to obtain the null distribution for the difference in mean wIBS values with each of the individuals in our sample, under the assumption of no difference between them. For a single test, each of the individuals in our sample was deemed to be more closely related to population A than B, when 5% or fewer values from the null distribution were greater than the observed (non-permuted) difference between A and B. These P values were further Bonferroni adjusted for the number of tests performed.

Figure 2D in the main text shows a heat map of the normalized mean wIBS scores. Marked with a black X are the populations that can potentially be excluded as source populations based on the permutation test. Figure S10 shows the individual plots for each of the STHs using the non-normalized values for wIBS.

**D-statistics**

We computed a four population test implemented as D-statistics (30) of the form *D(O, STH; refwIBS, REF)* where *O* is an outgroup, *STH* is each of the individuals, *refwIBS* is the population with the highest wIBS value for that individual, and *REF* is all of the other reference populations. We used the Tygray from Ethiopia as the outgroup. The D-statistics were computed using AdmixTools and the West Central Africa reference panel, used also for the wIBS analysis. This statistic measures whether the data is consistent with a four-population tree. In the tested framework *D(O, STH; refwIBS, REF), refwIBS* and *REF* form a clade with each other, to the exclusion of the other two (*STH* and the *Outgroup*). Under the assumption that the chosen *STH* sample has the highest genetic affinity to the *refwIBS* population (i.e. the population with the maximum wIBS statistic) we expect that the *D*-statistics computed under the tree shown above – *D(O, STH; refwIBS, REF)* – are always less than 0. Depending on the genetic similarity between the different reference populations, *D*-statistics might not be significantly less than 0 (Z-score <-3), but should not be significantly greater than 0 (Z-score >3). Our results, summarized in Figure S11, show that none of the non-*refwIBS* reference populations are significantly closer to the *STH* sample than the *refwIBS* population.

To further statistically test if the refwIBS populations have the highest genetic affinity with the corresponding *STH* individual, we performed a $X^2$ test for the proportion of D statistics that were above or below 0. If the two populations in the ingroup of the tree used for computing the D-statistic were randomly chosen, we would expect the proportion of D-statistics > 0 to be 0.5. For all STH individuals except one, the $X^2$ test showed that the distribution of D-statistics is due to chance with statistical significance of $p < 0.01$ (Table S11).

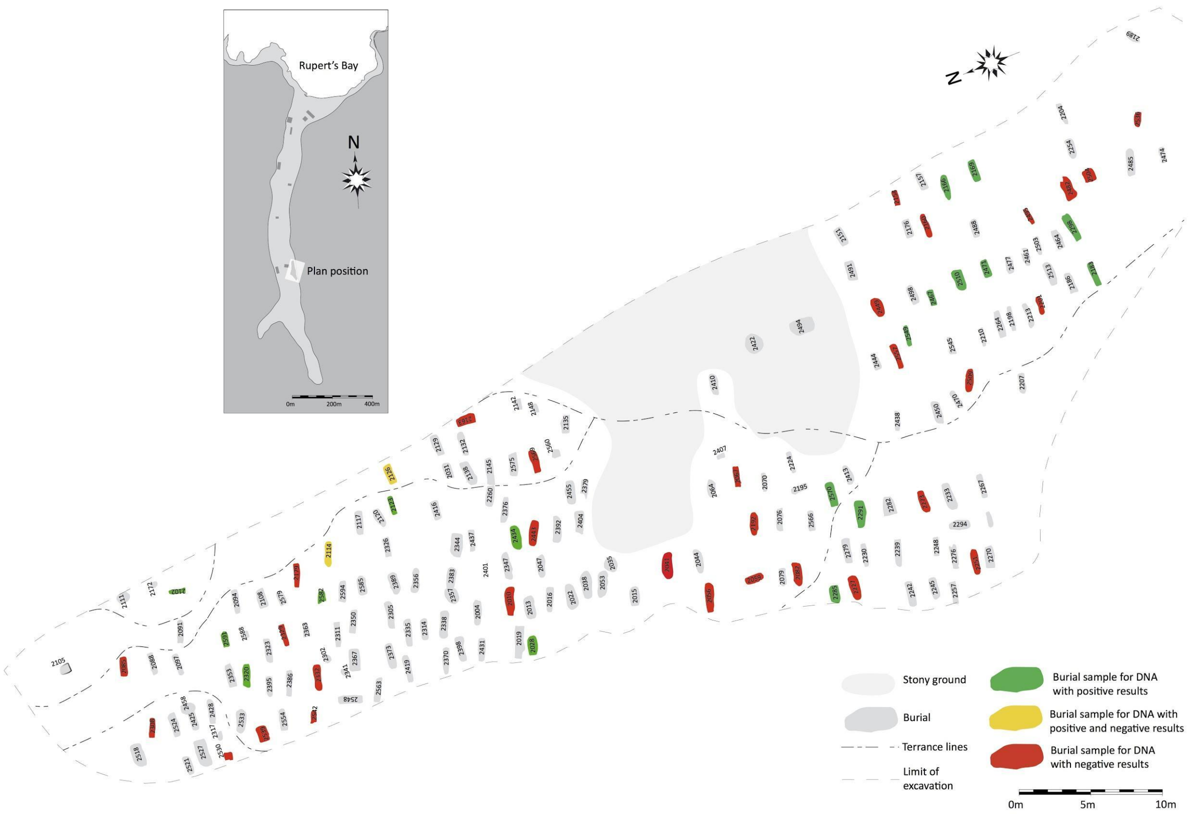


**Fig. S1.** Burial plan of the 178 burials containing 325 individuals excavated between 2007 and 2008, showing the location of the burials sampled for ancient DNA (marked in color) and the 20 burials from which low-coverage aDNA was recovered (marked in green and yellow). Burials in yellow correspond to multiple burials from which DNA could only be retrieved from one individual.

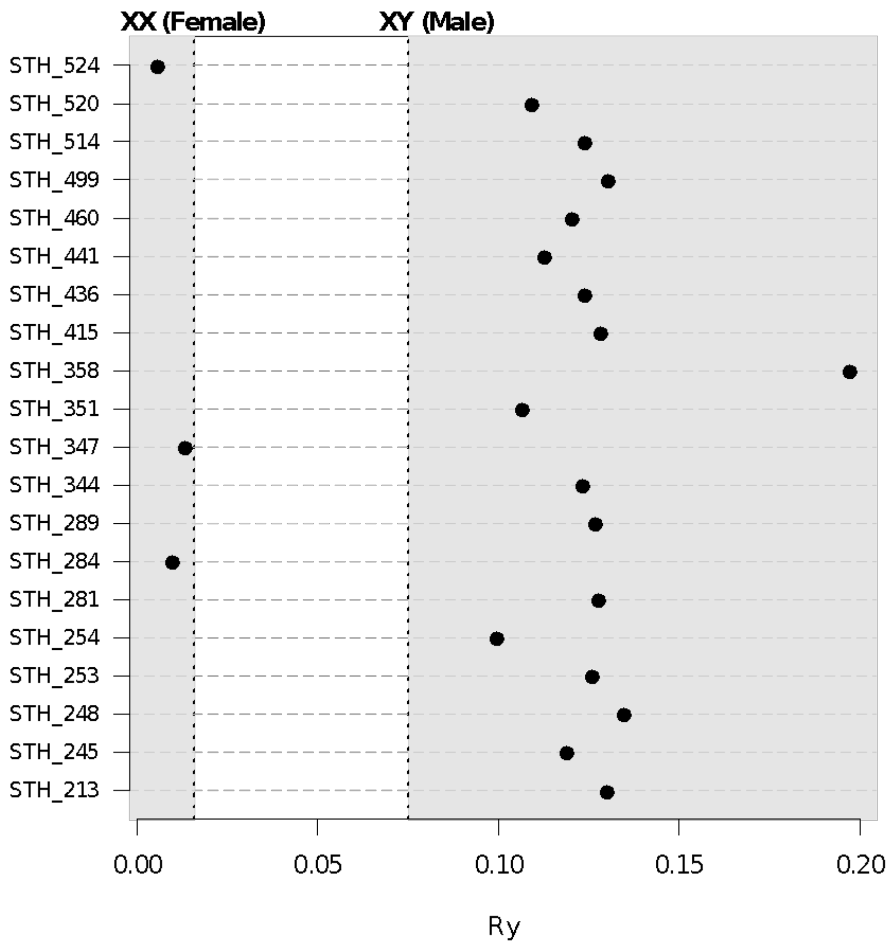


**Fig. S2.** Chromosomal sex of STH individuals. Ry is the ratio of reads aligning to the Y chromosome to those aligning to the X chromosome.

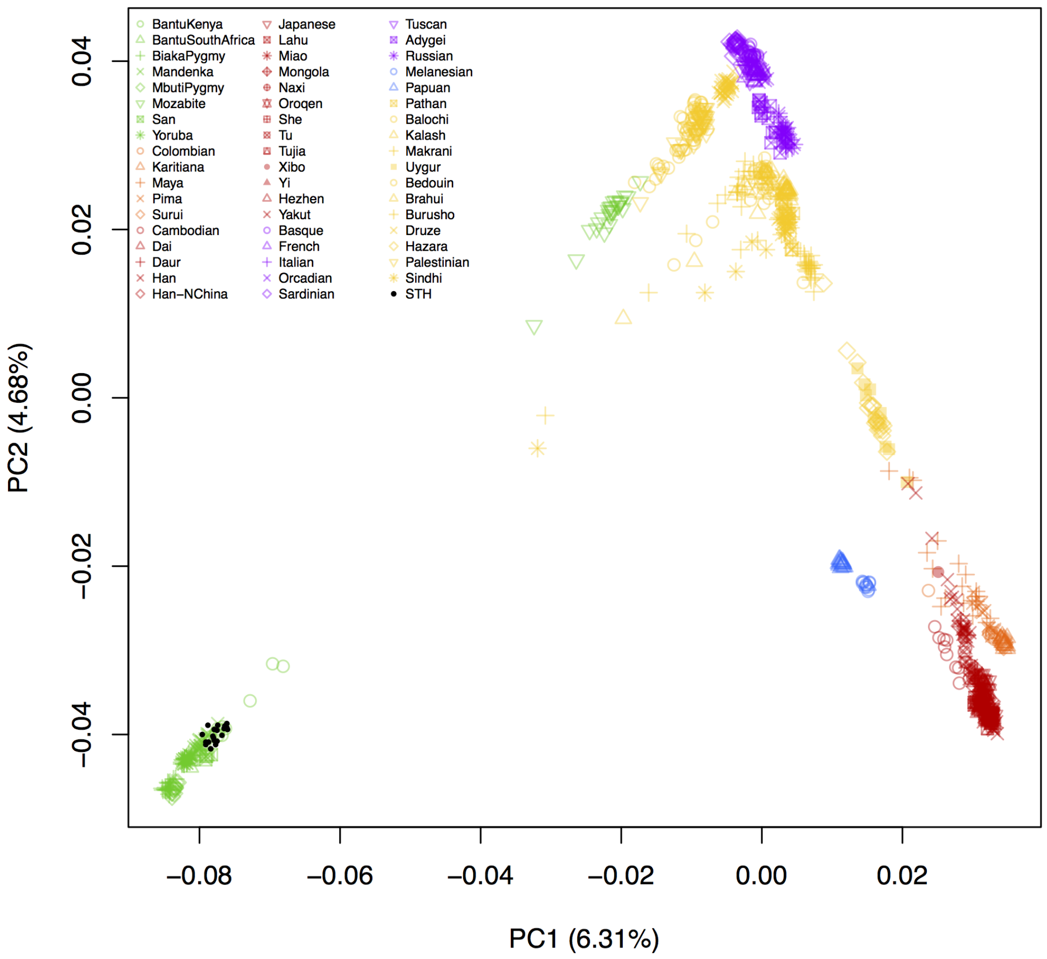


**Fig. S3.** Global PCA of overlapping sites using the publicly available reference panel HGDP (21, 22) based on 644,117 SNPs. The average number of overlapping sites between the panel and the samples is 103,093 SNPs (Table S9).

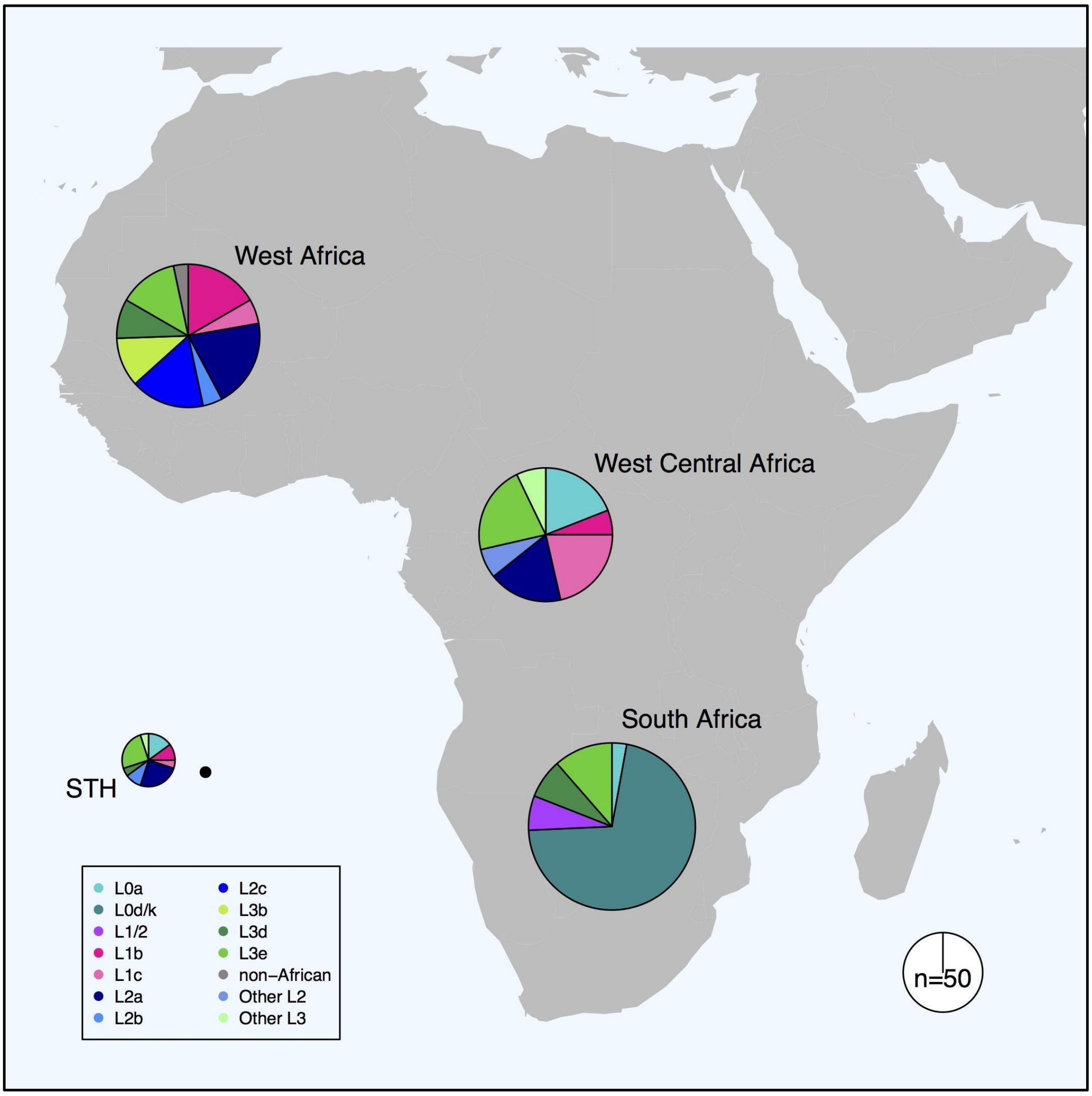


**Fig. S4.** Mitochondrial DNA haplogroup distributions for the STH individuals (n = 20) and different African regions (31).

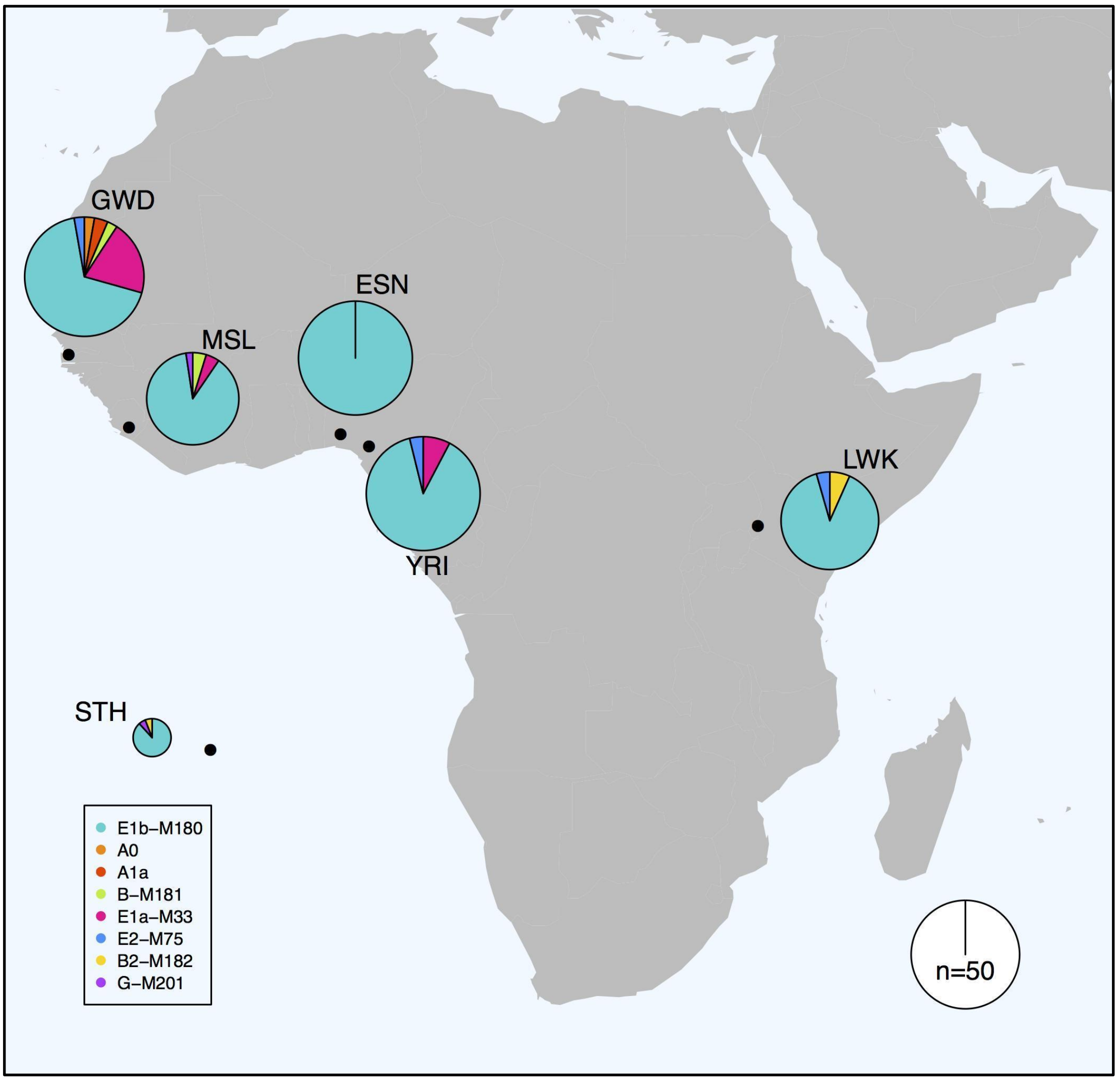


**Fig. S5.** Y-chromosome haplogroup distributions for the male STH individuals (n = 17) and different sub-Saharan reference populations (32).

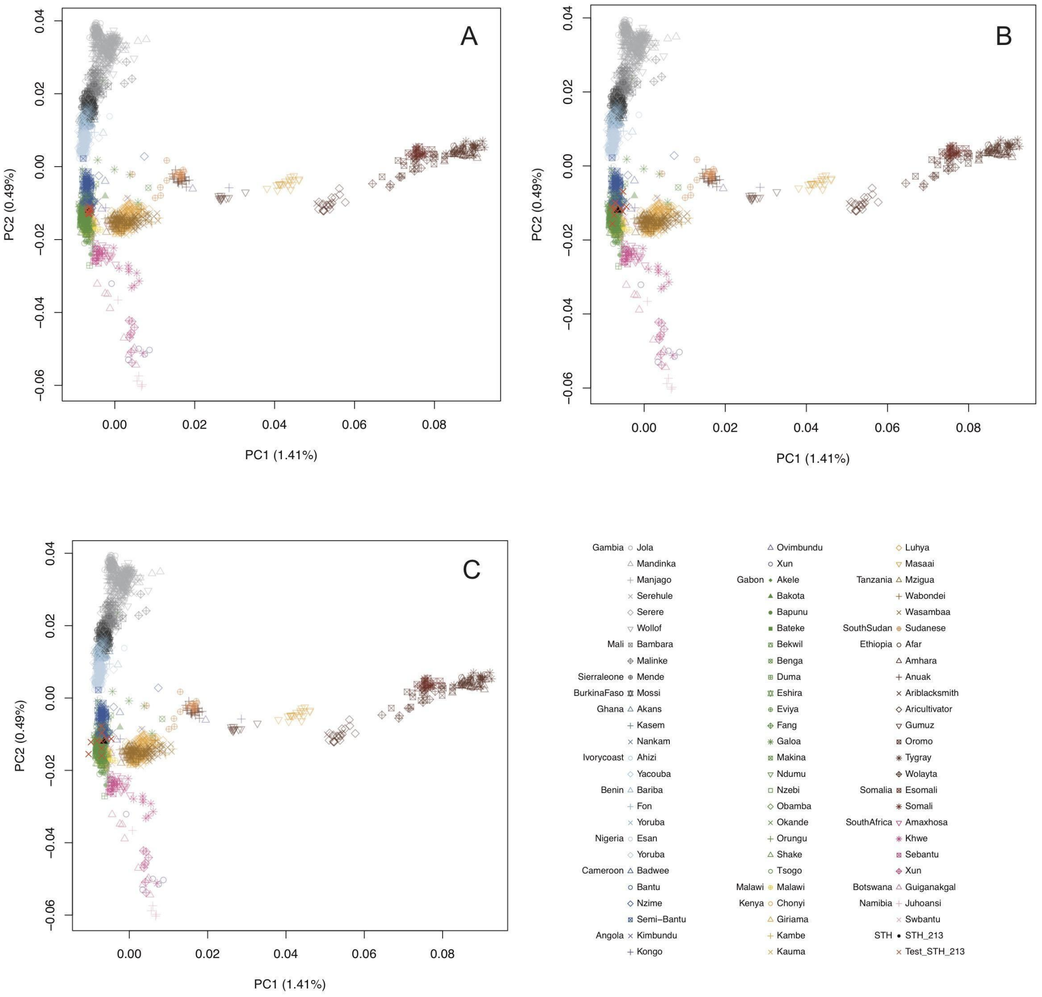


**Fig. S6.** Downsampling experiment of overlapping sites. PCA plots for individual STH 213 using 60k (**A**), 30k (**B**) and 15k (**C**) overlapping sites. The red Xs correspond to 10 iterations of the random sampling of overlapping sites.

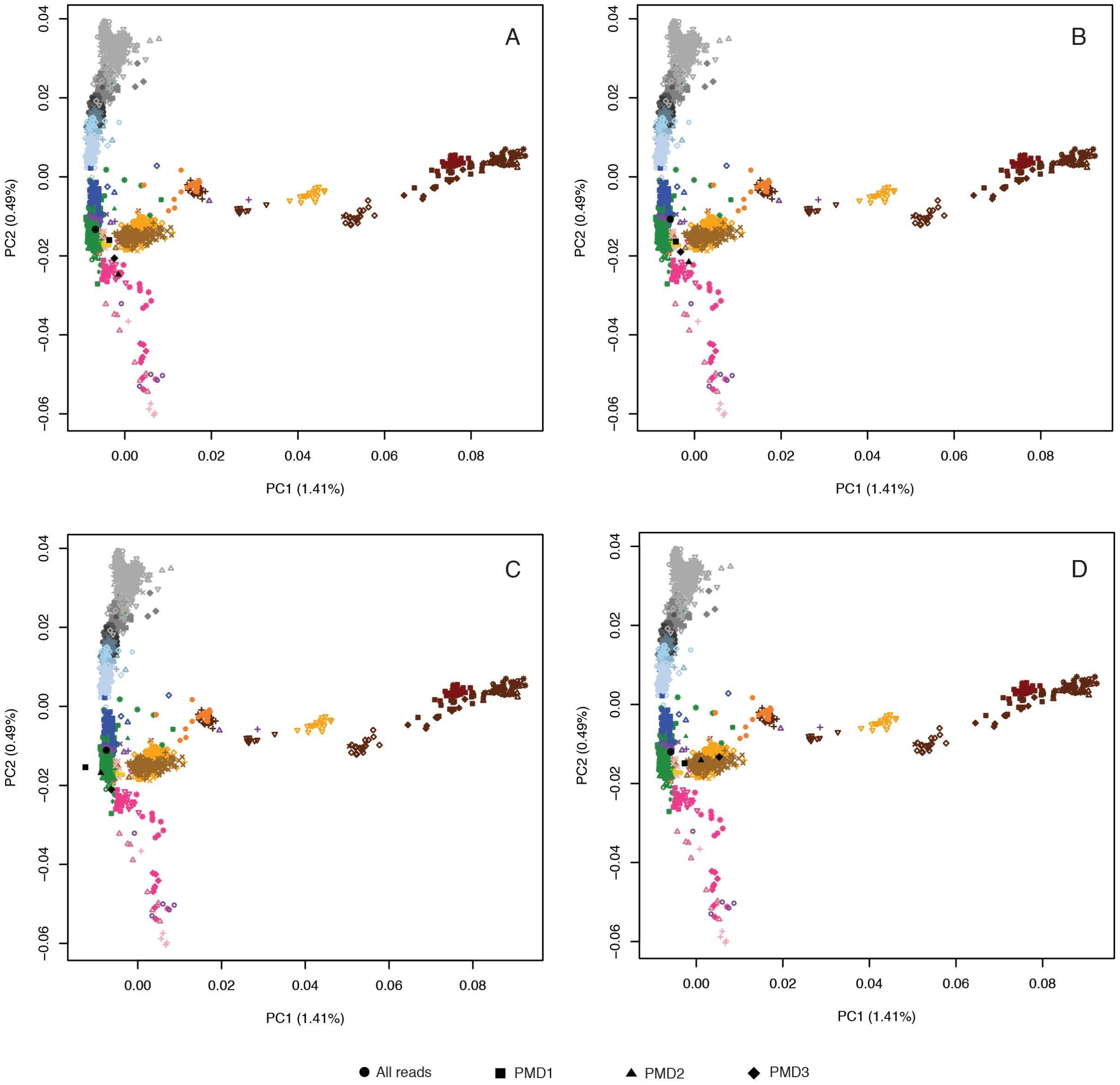


**Fig. S7.** PCA projection using all reads and only reads with signs of aDNA damage (PMD1, PMD2, PMD3) fom non-contaminated samples (STH 213 and STH 460, **A** and **B** respectively) and contaminated samples (STH 254 and STH 351, **C** and **D** respectively).

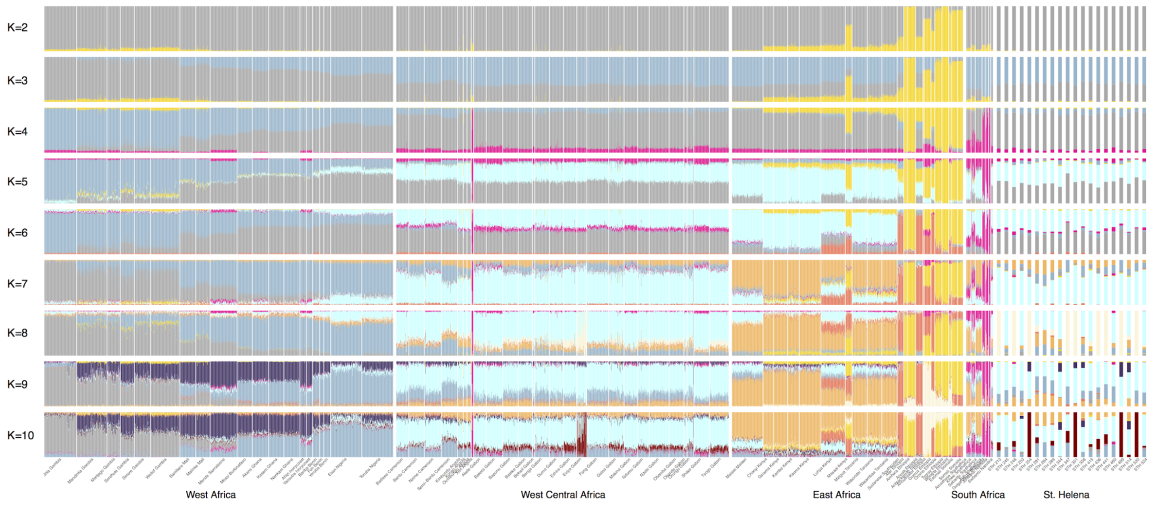


**Fig. S8.** Admixture analysis showing the ancestry proportions of the 20 STH individuals and the whole Africa reference panel comprising 2898 individuals from 76 different African populations spanning West West Central, South and East African regions. Plots were generated using a maximum-likelihood approach implemented in ADMIXTURE (28) and show the runs with the highest log-likelihood from K=2 to K=10.

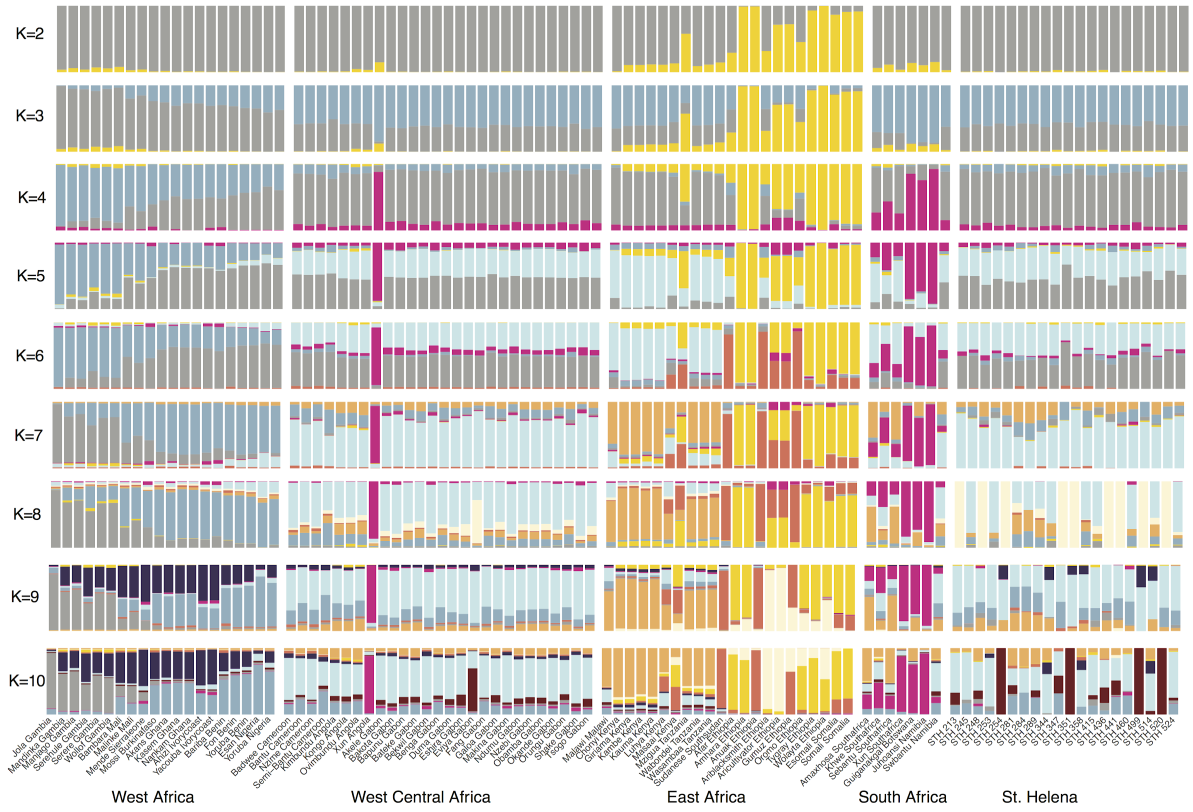


**Fig. S9.** Admixture analysis showing the ancestry proportions of the 20 STH individuals and the average ancestry proportions per population of the whole Africa reference panel comprising 2898 individuals from 76 different populations spanning West West Central, South and East African regions. Plots were generated using a maximum-likelihood approach implemented in ADMIXTURE (28) and show the runs with the highest log-likelihood from K=2 to K=10.

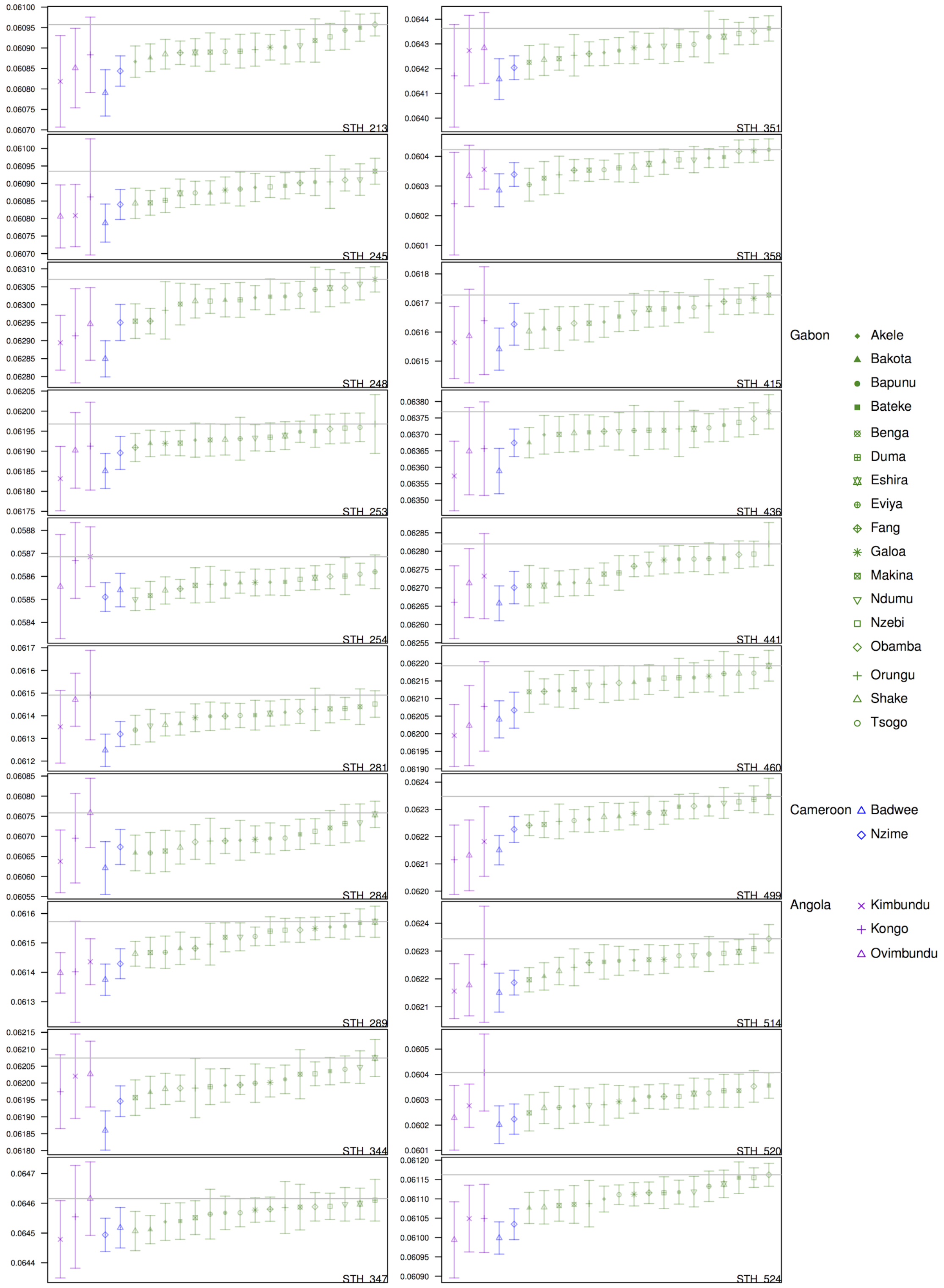


**Fig. S10.** Plots of non-normalized mean wIBS scores. The horizontal line corresponds to the highest wIBS value and the symbol that intersects the line corresponds to the population with the highest wIBS score, namely the one suggested as being the more closely related. The populations which standard-deviation values also intersect the line can't be excluded with confidence as being also related.

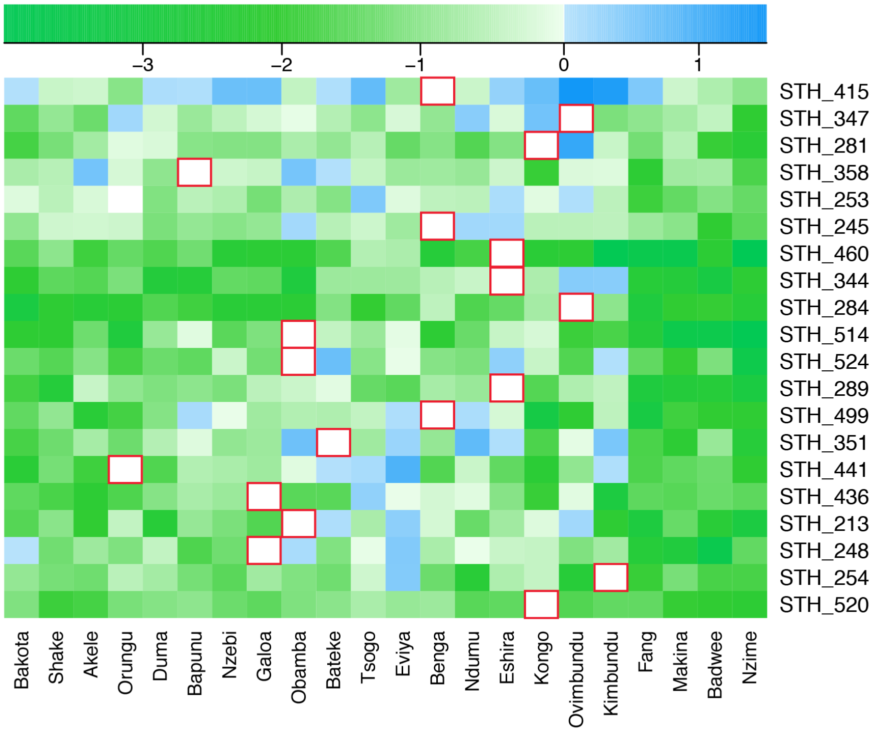


**Fig. S11.** Heat map of Z-scores showing in red squares the more closely related population for each of the STH individuals according to the wIBS statistic.

### References


1. Pearson AF (2016) *Distant Freedom: St Helena and the Abolition of the Slave Trade, 1840-1872* (Liverpool University Press).

2. Pearson AF, Jeffs B, Witkin A, MacQuarrie H (2011) *Infernal traffic: Excavation of a liberated African graveyard in Rupert's Valley, St. Helena* (Council for British Archeology).

3. Dabney J, et al. (2013) Complete mitochondrial genome sequence of a Middle Pleistocene cave bear reconstructed from ultrashort DNA fragments. *Proc Natl Acad Sci U S A* 110(39):15758–15763.

4. Rohland N, Hofreiter M (2007) Ancient DNA extraction from bones and teeth. *Nat Protoc* 2(7):1756–1762.

5. Meyer M, Kircher M (2010) Illumina Sequencing Library Preparation for Highly Multiplexed Target Capture and Sequencing. *Cold Spring Harbor Protocols* 2010(6):db.prot5448–pdb.prot5448.

6. Lindgreen S (2012) AdapterRemoval: easy cleaning of next-generation sequencing reads. *BMC Res Notes* 5:337.

7. Li H, Durbin R (2009) Fast and accurate short read alignment with Burrows–Wheeler transform. *Bioinformatics* 25(14):1754–1760.

8. Andrews RM, et al. (1999) Reanalysis and revision of the Cambridge reference sequence for human mitochondrial DNA. *Nat Genet* 23(2):147.

9. Li H, et al. (2009) The Sequence Alignment/Map format and SAMtools. *Bioinformatics* 25(16):2078–2079.

10. Jónsson H, Ginolhac A, Schubert M, Johnson PLF, Orlando L (2013) mapDamage2.0: fast approximate Bayesian estimates of ancient DNA damage parameters. *Bioinformatics* 29(13):1682–1684.

11. Briggs AW, et al. (2007) Patterns of damage in genomic DNA sequences from a Neandertal. *Proc Natl Acad Sci U S A* 104(37):14616–14621.

12. Skoglund P, et al. (2014) Separating endogenous ancient DNA from modern day contamination in a Siberian Neandertal. *Proc Natl Acad Sci U S A* 111(6):2229–2234.

13. Renaud G, Slon V, Duggan AT, Kelso J (2015) Schmutzi: estimation of contamination and endogenous mitochondrial consensus calling for ancient DNA. *Genome Biol* 16:224.

14. Fu Q, et al. (2013) A revised timescale for human evolution based on ancient mitochondrial genomes. *Curr Biol* 23(7):553–559.

15. Green RE, et al. (2008) A complete Neandertal mitochondrial genome sequence determined by high-throughput sequencing. *Cell* 134(3):416–426.

16. Skoglund P, Storå J, Götherström A, Jakobsson M (2013) Accurate sex identification of ancient human remains using DNA shotgun sequencing. *J Archaeol Sci* 40(12):4477–

4482.

17. Monroy Kuhn JM, Jakobsson M, Günther T (2018) Estimating genetic kin relationships in prehistoric populations. *PLoS One* 13(4):e0195491.

18. Korneliussen TS, Albrechtsen A, Nielsen R (2014) ANGSD: Analysis of Next Generation Sequencing Data. *BMC Bioinformatics* 15:356.

19. Cruz-Dávalos DI, et al. (2018) In-solution Y-chromosome capture-enrichment on ancient DNA libraries. *BMC Genomics* 19(1). doi:10.1186/s12864-018-4945-x.

20. Poznik GD, et al. (2016) Punctuated bursts in human male demography inferred from 1,244 worldwide Y-chromosome sequences. *Nat Genet* 48(6):593–599.

21. Li JZ, et al. (2008) Worldwide human relationships inferred from genome-wide patterns of variation. *Science* 319(5866):1100–1104.

22. Malaspinas A-S, et al. (2014) bammds: a tool for assessing the ancestry of low-depth whole-genome data using multidimensional scaling (MDS). *Bioinformatics* 30(20):2962–2964.

23. Patin E, et al. (2014) The impact of agricultural emergence on the genetic history of African rainforest hunter-gatherers and agriculturalists. *Nat Commun* 5:3163.

24. Patin E, et al. (2017) Dispersals and genetic adaptation of Bantu-speaking populations in Africa and North America. *Science* 356(6337):543–546.

25. Patterson N, Price AL, Reich D (2006) Population Structure and Eigenanalysis. *PLoS Genetics* 2(12):e190.

26. Skoglund P, et al. (2012) Origins and genetic legacy of Neolithic farmers and hunter-gatherers in Europe. *Science* 336(6080):466–469.

27. Purcell S, et al. (2007) PLINK: a tool set for whole-genome association and population-based linkage analyses. *Am J Hum Genet* 81(3):559–575.

28. Alexander DH, Novembre J, Lange K (2009) Fast model-based estimation of ancestry in unrelated individuals. *Genome Res* 19(9):1655–1664.

29. Jagadeesan A, et al. (2018) Reconstructing an African haploid genome from the 18th century. *Nat Genet* 50(2):199–205.

30. Patterson N, et al. (2012) Ancient admixture in human history. *Genetics* 192(3):1065–1093.

31. Salas A, et al. (2002) The making of the African mtDNA landscape. *Am J Hum Genet* 71(5):1082–1111.

32. Cruciani F, et al. (2011) REPOR TA Revised Root for the Human Y Chromosomal Phylogenetic Tree: The Origin of Patrilineal Diversity in Africa. *Am J Hum Genet* 88(6):814–818.
